# CLOCK-dependent pathway in a single pair of LN_d_ neurons instruct circadian-independent interval timing behavior

**DOI:** 10.1101/2025.09.17.676708

**Authors:** Hongyu Miao, Zekun Wu, Yanan Wei, Woo Jae Kim

## Abstract

Interval timing is a cognitive ability essential for behaviors such as mating, foraging, and decision-making, and it is distinct from circadian rhythm regulation. Despite the involvement of circadian clock genes in both interval timing and circadian rhythms, the mechanisms differentiating these functions remain unclear. Using *Drosophila* as a model, we demonstrate that the CLK/CYC heterodimer, but not PER/TIM, is essential for interval timing. Neuronal CLK/CYC expression is necessary and sufficient for sexual experience-dependent shorter mating duration (SMD) behavior. We identified that CLK/CYC expression in a single pair of ITP-positive LN_d_ neurons is pivotal for SMD. These neurons are glutamatergic with output circuits to central brain regions. CLK variants lacking DNA binding motifs dissociate circadian rhythms from interval timing and sleep behaviors in these neurons. Our study uncovers a specialized circuit for interval timing and highlights the non-circadian functions of circadian clock genes.

**Statement of Significance:** This study in *Drosophila* demonstrates that the CLK/CYC heterodimer is crucial for interval timing, distinct from circadian rhythm regulation. The research pinpoints a specific pair of neurons that are critical for shortened mating duration behavior, and it suggests that CLK variants can influence interval timing dissociation from circadian rhythm and sleep behaviors.

**Highlights:**

- CLK/CYC heterodimer are essential for interval timing.
- Specialized mechanism of interval timing behavior revealed.
- Identified a pair of glutamatergic ITP-LN_d_ neurons that independently regulate both interval timing and sleep.
- Revealed a novel mechanism by which CLK variants regulate sleep and interval timing.

**Graphical Abstract:** Different CLK/CYC mediate separate regulatory mechanism between circadian rhythm, sleep and interval timing in *Drosophila melanogaster*.

## Introduction

The precise measurement of time intervals, ranging from seconds to minutes, is a critical cognitive function that is essential for behaviors such as mating, foraging, and navigation^1–3^. Interval timing is thought to encompass an internal clock mechanism that depends on the synchronization of pacemaker-accumulator circuits with memory circuits in the brain^4–7^. *Drosophila melanogaster*, with its well-characterized neural circuits and molecular mechanisms, serves as an excellent model organism for the study of interval timing^8^. Interval timing has been identified in the mating behaviors of *Drosophila*, particularly in the duration of male mating. Males demonstrate the ability to adjust their mating duration based on contextual experiences, suggesting the formation of long-term memory to optimize their sexual investment^9–13^.

The mating duration of male fruit flies serves as an excellent model for investigating interval timing, as it is influenced by both internal states and environmental contexts. Previous research by our group ^9–11,14–16^ and others ^17–19^ has established robust frameworks for studying mating duration using advanced genetic tools, enabling the dissection of neural circuits underlying interval timing. Notably, males exhibit prolonged mating duration when exposed to rival environments ^9,10,20^. In contrast, they display shortened mating duration (SMD) behavior under sexually saturated conditions, where they reduce their mating investment ^11^. These findings highlight the adaptability of mating behavior in response to social and environmental cues, providing a valuable system for exploring the neural and molecular mechanisms of interval timing.

The circadian rhythm in *Drosophila melanogaster* has been a subject of intense research for several decades, and significant progress has been made in understanding the molecular and genetic underpinnings of this biological clock^21–23^. Key genes such as *period (per)*, *timeless (tim)*, *Clock (Clk)* and *cycle (cyc)* have been identified and characterized, providing insights into the feedback loops that regulate the circadian rhythm at the molecular level^24^. These genes form the core of the molecular clock, with their timed expression, localization, post-transcriptional modification, and function being critical for maintaining the circadian cycle. Various regulators, including phosphatases and kinases, act on different steps of this feedback loop to ensure strong and accurately timed rhythms^25–27^.

Circadian clock plays a role in regulating sleep by ensuring that it occurs at the appropriate time. However, the quantity and quality of sleep are also influenced by other systems that maintain a balance between sleep and wakefulness. Sleep is regulated by separate genetic and cellular mechanisms that control the need for sleep, adjust to environmental signals, and react to extended periods of alertness. In addition, the regulation of sleep according to the body’s internal clock requires multiple groups of cells and molecules that coordinate sleep and waking cycles at precise times of the day, rather than being governed by a single oscillating signal^28,29^.

Circadian rhythm pertains to the 24-hour cycle that governs biological processes, whereas sleep is a reversible state characterized by diminished activity and responsiveness. Interval timing refers to the measurement of durations and intervals, such as the mating duration or foraging actions^20,30,31^. The primary distinction lies in the periodicity of circadian rhythms and sleep, which adhere to fixed daily patterns, while interval timing is adaptable and not constrained by a 24-hour cycle, enabling organisms to measure time intervals according to their specific requirements^32^.

Although the connection between circadian timing and different physiological processes has been extensively studied, the genetic understanding of the interaction between circadian timing and interval timing remains incomplete. Our previous research demonstrated that circadian clock genes, specifically *per* and *tim*, rather than *Clk* and *cyc*, modulate the rival-induced longer-mating-duration (LMD) behavior, a distinct form of interval timing critical for maximizing sperm competition^9,10,20^.

In this study, we investigate the role of the *Clk*/*cyc* gene complex and associated factors within a single pair of LN_d_ neurons in modulating the sexually experienced-dependent shorter-mating-duration (SMD) behavior, which has been previously shown to be elicited by gustatory and pheromonal cues from females^11^.

## Results

### Neuronal expression of CLK/CYC heterodimer is necessary and sufficient to generate sexual experience-mediated reduced mating investment

In the context of core circadian rhythm genes, it was observed that mutations in the *per* or *tim* genes, as well as compound mutations in both *per* and *tim*, did not disrupt SMD behavior (Fig. 1A-C). However, mutations in the *Clk* and *cyc* genes resulted in the absence of SMD behavior (Fig. 1E-F). Additionally, mutations in the *cryptochrome* (*cry*) gene (*cry^03^*) and the sleepless mutant of *quiver* (*qvr^01^*)^33^ did not alter the SMD behavior (Fig. 1D and Fig. S1A). These findings suggest that the CLK/CYC heterodimer is uniquely involved in interval timing behaviors among the core circadian rhythm gene components.

**Figure 1.**
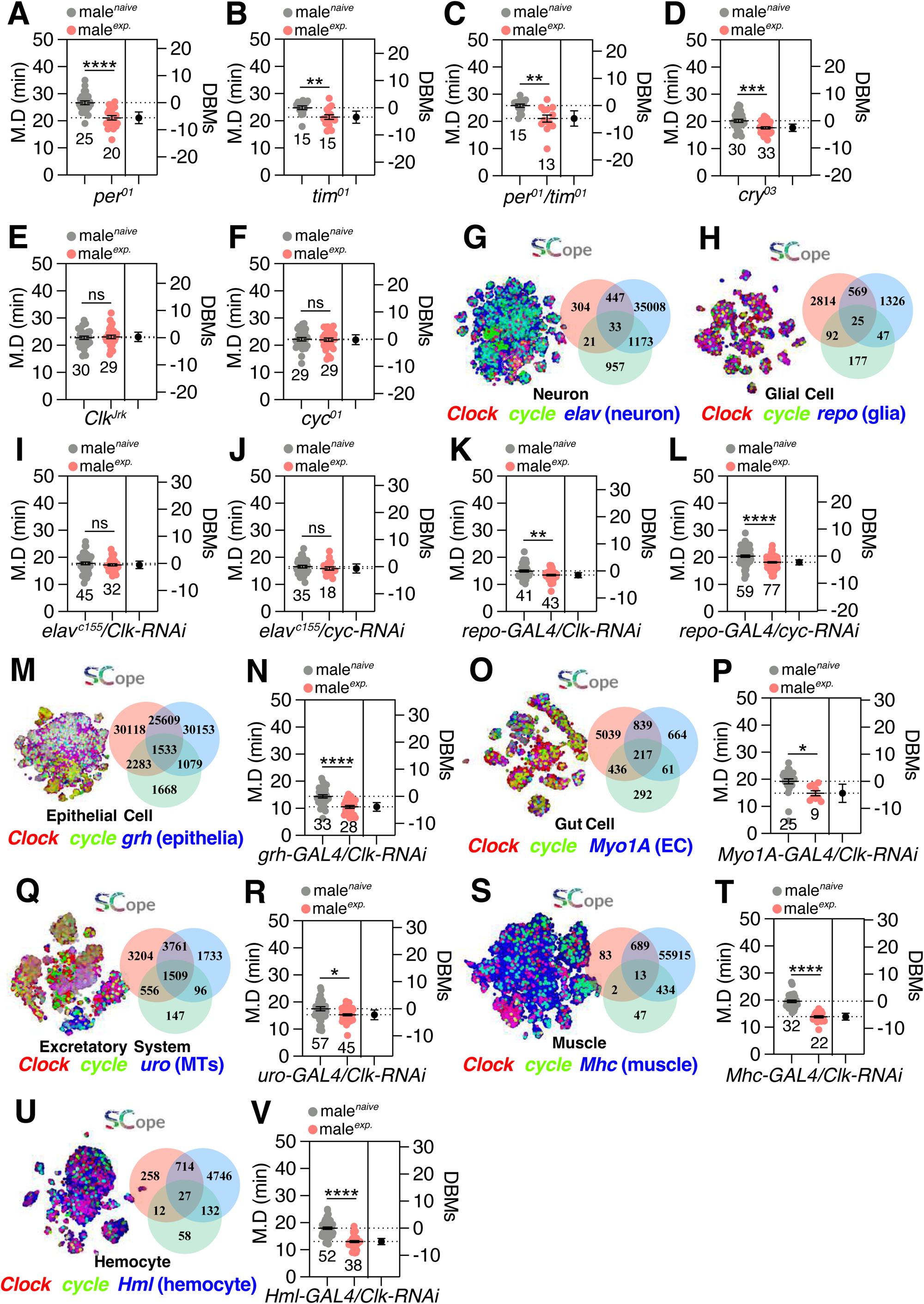
Interval timing is regulated by neuronal expression of CLK/CYC heterodimer, not PER/TIM. (A-F) SMD assays of core circadian rhythm gene mutations. Light gray dots represent naïve males, and pink dots represent experienced ones. Dot plots represent the M. D (mating duration) of each male fly. The mean value and standard error are labeled within the dot plot (black lines). Asterisks represent significant differences, as revealed by the Student’s t test, and ns represents non-significant differences (*\*p<0.05, **p<0.01, ***p< 0.001, ****p< 0.0001*). DBMs refers to the Difference between Means. For detailed methods, see the “Methods” for a detailed description of the mating duration assay used in this study. In the framework of our investigation, the routine application of internal controls is employed for the vast majority of experimental procedures, as delineated in the “**Mating Duration Assay**” and “**Statistical Tests**” subsections of the Methods section. The numerical values in the bar graph denote the number of fruit flies in the respective experimental groups, referred to as “n”. Subsequent estimated statistical graphs will employ the same numerical designations. (G-H) Fly SCope single-cell RNA sequencing data of cells co-expressing *Clk/cyc* together with (G) *elav* in the ‘Neuron’ population and (H) *repo* in the ‘Glial Cell’ population.. Annotations and gene names of all the above data are color-coded using red, green, and blue words. When cells overlap, the color of the dots is either yellow, cyan, or magenta. The Venn diagram indicates the number of cells in each overlap category. (I-J) SMD assays for *elav^c155^*-mediated neuronal knockdown of *Clk/cyc* via *Clk-RNAi* and *cyc-RNAi*. (K-L) SMD assays for *repo-GAL4*-mediated glial knockdown of *Clk/cyc* via *Clk-RNAi* and *cyc-RNAi*. (M-V) SCope data of *Clk/cyc* expression in different tissue populations and SMD assays for tissue-specific knockdown of *Clk* via *Clk-RNAi* using (N) *grh-GAL4*, (P) *Myo1A-GAL4*, (R) *uro-GAL4*, (T) *Mhc-GAL4* and (V) *Hml-GAL4*.

The targeted RNAi-mediated knockdown of *Clk* or *cyc* in neurons resulted in the disruption of SMD behavior (Fig. 1I-J and Fig. S1B-C). Additionally, analysis of the fly RNAseq dataset platform, fly SCope, predicted co-expression of *Clk* and *cyc* in specific neuronal populations (Fig. 1G)^34^. While co-expression of *Clk* and *cyc* in glial cells was observed (Fig. 1H), knockdown of these genes in glial populations did not affect SMD behavior (Fig. 1K-L), indicating that glial expression of CLK/CYC is not essential for interval timing behavior. Furthermore, knockdown of *Clk* in epithelial, gut, Malpighian tubules (MTs), muscle, hemocyte, and intestinal stem cells (ISCs), despite their high expression of *Clk* and *cyc*, did not alter SMD behavior (Fig. 1M-V and Fig. S1K-L). Collectively, the genetic control experiments and the testing of independent RNAi lines suggest that only neuronal expression of *Clk* and *cyc* is required for the sexual experience-mediated reduction in mating investment (Fig. S1D-J). These findings underscore the specificity of the neuronal CLK/CYC expression in mediating the behavioral changes associated with sexual experience, highlighting the importance of cell-type-specific gene expression in the regulation of complex behaviors.

### The expression of CLK within NPF-expressing cry-positive circadian neurons in the brain is critical for interval timing behavior

*Clk* and *cyc* are robustly expressed in neuronal populations throughout the *Drosophila* body (Fig. 2A). However, targeted knockdown of *Clk* in all neurons except those of the ventral nerve cord (VNC) still resulted in the disruption of SMD behavior (Fig. 2B), indicating that *Clk* expression within the VNC is not essential for the generation of SMD behavior. Furthermore, the selective knockdown of *Clk* or *cyc* in the brain alone was sufficient to impair SMD behavior (Fig. 2C-E), suggesting that among the diverse neuronal populations, only CLK/CYC expression within the brain is necessary for the manifestation of SMD behavior.

**Figure 2.**
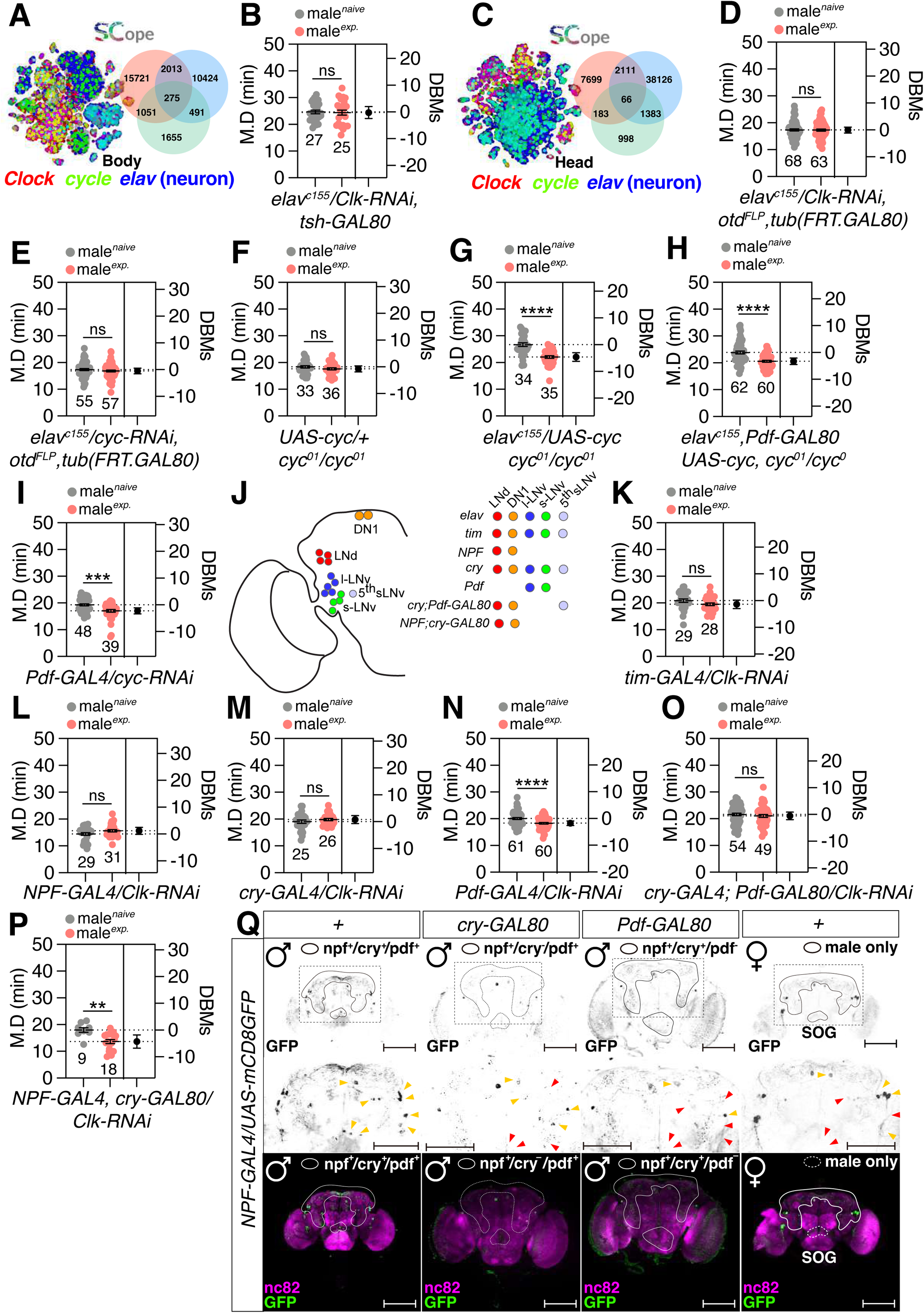
NPF and CRY co-expressed clock neurons in the brain are critical for the generation of interval timing behavior. (A) Fly SCope single-cell RNA sequencing data of cells co-expressing *Clk/cyc* together with *elav* in ‘Body’ population. (B) SMD assay for *elav^c155^* driver mediated knockdown of *Clk* in tsh-negative neurons via *Clk-RNAi; tsh-GAL80.* (C) Fly SCope single-cell RNA sequencing data of cells co-expressing *Clk/cyc* together with *elav* in ‘Head’ population. (D-E) SMD assays for *elav^c155^* drivers mediated knockdown of *Clk/cyc* in head neurons via (D) *tub(FRT.GAL80); otd^FLP^; Clk-RNAi* and (D) *tub(FRT.GAL80); otd^FLP^; cyc-RNAi.* (F-H) Genetic rescue experiments of SMD assays for GAL4 mediated overexpression of CYC via (G) *elav^c155^*(H) *elav^c155^; Pdf-GAL80* in *cyc* mutant background flies. (I) SMD assays for *Pdf-GAL4*-mediated knockdown of *Clk* in core oscillator cells via *Clk-RNAi.* (J) Diagram of circadian-associated driver-labeled clock cells. Subsets of neurons labeled by GAL4 drivers named in italics are color-coded. (K-P) SMD assays for circadian-related-GAL4 mediated knockdown of *Clk* via *Clk-RNAi* using the (K) *tim-GAL4*, (L) *NPF-GAL4*, (M) *cry-GAL4*, (N) *Pdf-GAL4*, (O) *cry-GAL4; Pdf-GAL80*, and (P) *NPF-GAL4; Pdf-GAL80*. (Q) Male (left three) and female (right one) flies brain expressing *npf-GAL4; UAS-mCD8GFP* together with *cry-GAL80* and *Pdf-GAL80* were immunostained with anti-GFP (green) and nc82 (magenta) antibodies. The white circles denote neurons that express various genes, with the genotypes specified to the right of the circles. The yellow arrow indicates NPF-positive cells, whereas the red arrow represents NPF-positive cells that are absent in this genotype. Scale bars represent 50 μm. For detailed methods, see the “Methods” for a detailed description of the immunostaining procedure used in this study.

Through a genetic rescue approach targeting the *cyc* gene, we discovered that neuronal expression of *cyc*, with the exception of the PDF-expressing lateral ventral neurons (LN_v_), was sufficient to restore SMD behavior (Fig. 2F-H), despite the rescue’s inability to restore circadian rhythmicity^35^. Additionally, the knockdown of *cyc* specifically in PDF neurons did not affect SMD behavior (Fig. 2I), implying that CLK/CYC expression in core oscillator cells, such as sLN_v_ and lLN_v_, is not required for the generation of interval timing behavior.

Utilizing a suite of GAL4 drivers specific to clock cells in conjunction with *Clk-RNAi* (Fig. 2J and Fig. S2A), we conducted a detailed analysis revealing that the expression of *Clk* and *cyc* in neurons positive for neuropeptide F (NPF) and cry is essential for the generation of SMD behavior (Fig. 2K-O and Fig. S2B-C). Knockdown of *Clk* in neurons that are NPF-positive but cry-negative did not impair SMD behavior (Fig. 2P), indicating that the expression of *Clk* in both cry-positive and NPF-positive neurons is critical for the manifestation of SMD behavior. NPF-expressing cry-positive neurons are located in the LN_d_ and DN regions of both male and female brains (Fig. 2Q). However, a unique male brain phenotype was observed, with NPF-positive and CRY-positive neuronal processes extending near the suboesophageal ganglion (SOG) region (Fig. 2Q), which is known to be important for processing taste information. Previous research from our laboratory has demonstrated the presence of male-specific CRY-positive and NPF-positive LN_d_ neurons^36^, and the role of these neurons’ sexual dimorphism in mating behavior has been reported^37,38^. These findings suggest that the sexual dimorphism in the distribution of NPF-expressing cry-positive neurons may underlie sex-specific differences in the processing of sensory information related to SMD behavior. Our results collectively identify a selective requirement for CLK/CYC within specific clock neurons in the modulation of SMD behavior.

### Two pair of clock neurons are associated with CLK function to generate interval timing

Utilizing the recently published transcriptomic taxonomy dataset for *Drosophila* circadian neurons^39^, we conducted a targeted screen to identify the minimal subset of Clk-expressing neurons necessary for the generation of SMD (Table. S1). Our analysis revealed that CLK expression within the *GAL4^R54D11^*-labeled ITP-LN_d_ and 5^th^ sLN_v_ neurons are pivotal for interval timing behavior (Table S1, Fig. 3A and S3A). Knockdown of *Clk* specifically in adults or excluding of the VNC using the *GAL4^R54D11^* driver was sufficient to impair SMD (Fig. 3B-D and Fig. S3B), indicating that adult brain expression of *Clk* in ITP-LN_d_ and 5^th^ sLN_v_ neurons are required to generate SMD behavior. No sexual dimorphism was observed in the expression pattern of the *GAL4^R54D11^*driver within the brain (Fig. S3C-D). Furthermore, the disruption of SMD following Clk knockdown with the *ITP^T2A^-GAL4* and LN_d_ drivers (*Mai179-GAL4*, marking ITP-LN_d_, 5^th^-sLN_v_, and 2 sNPF-LN_d_ neurons) confirmed that the *GAL4^R54D11^* driver-labelled neurons are indeed ITP-LN_d_ and 5^th^ sLN_v_ neurons (Fig. 3E-F).

**Figure 3.**
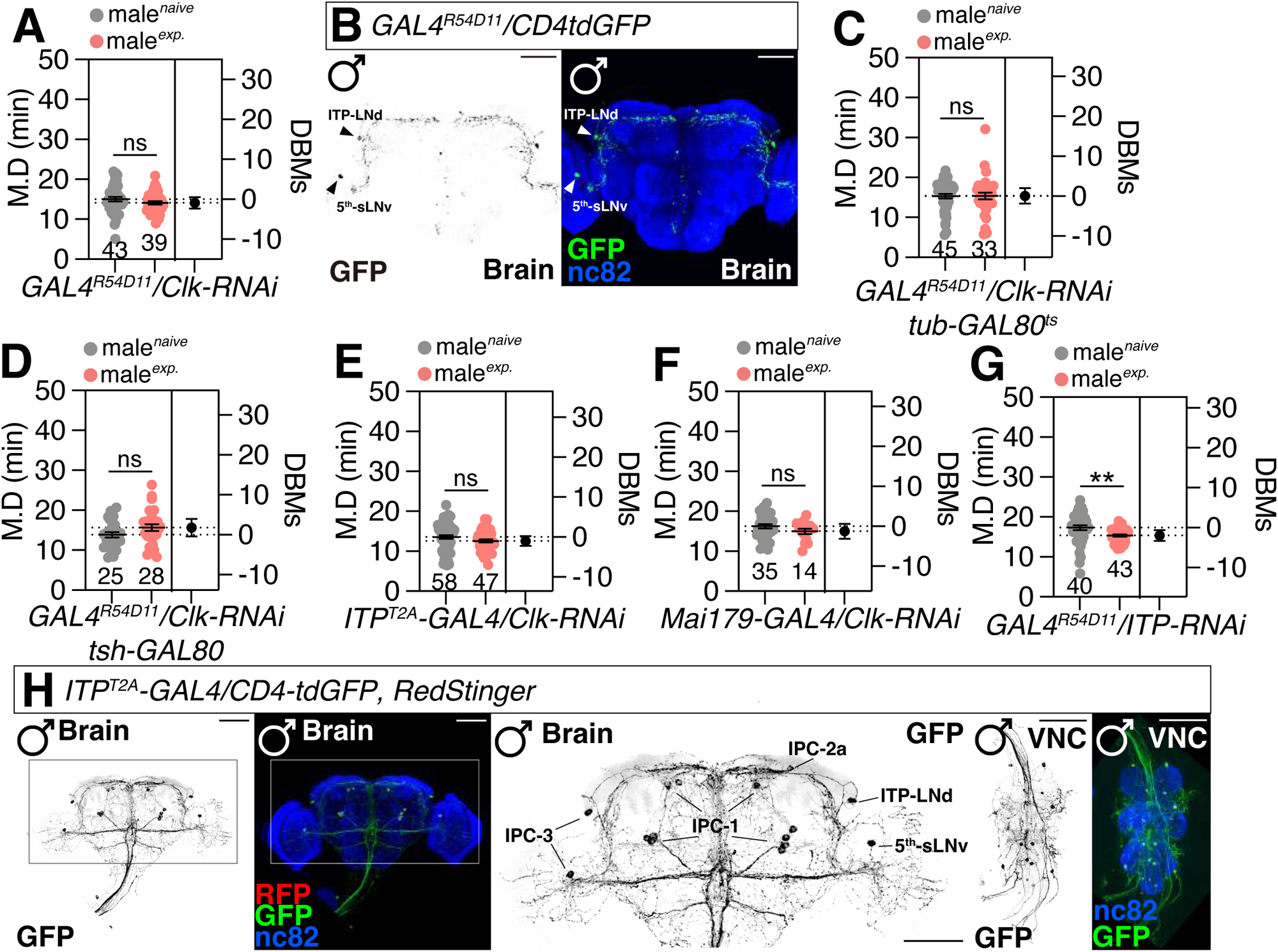
ITP-LN_d_ and 5^th^ sLN_v_ generate SMD behavior through the CLK function. (A) SMD assay for *GAL4^R54D11^* driver mediated knockdown of *Clk* via *Clk-RNAi*. (B) Brain of male flies expressing *GAL4^R54D11^* together with *UAS-CD4tdGFP* was immunostained with anti-GFP (green), and nc82 (blue) antibodies. Scale bars represent 50 μm. The left panel is presented as a gray scale to clearly show the cellular morphology in the adult brain labeled by *GAL4^R54D1^* driver. Yellow arrows indicate ITP-LN_d_ and 5^th^ sLN_v_ soma. (C) SMD assay for *GAL4^R54D11^*driver mediated adult-stage knockdown of *Clk* via *Clk-RNAi* together with *tub-GAL80^ts^*. After hybridization, parentals were placed at 22 °C, and after adults had eclosion, they were transferred to 29 °C and left for 5 days before the experiment. (D) SMD assay for *GAL4^R54D11^* driver mediated knockdown of *Clk* in tsh-negative neurons via *Clk-RNAi; tsh-GAL80*. (E-F) SMD assays for ITP-LN_d_ and 5^th^ sLN_v_ labeled-GAL4 mediated knockdown of *Clk* via *Clk-RNAi* using (E) *ITP^T2A^-GAL4* and (F) *Mai179-GAL4*. (G) SMD assay for *GAL4^R54D11^* driver mediated knockdown of *ITP* via *ITP-RNAi*. (H) Male flies brain and VNC expressing *ITP^T2A^-GAL4* driver together with *UAS-CD4tdGFP* and *UAS-RedStinger* were immunostained with anti-GFP (green), anti-DsRed (red), and anti-nc82 (blue) antibodies. Scale bars represent 50 μm in brain panels and 50 μm in VNC panels. Boxes indicate the magnified regions of interest presented in the middle panels. The text indicates the name of the *ITP^T2A^-GAL4*-labeled brain cells. IPC, ITP-producing cells.

ITP is a key endocrine regulator of water homeostasis in *Drosophila*^40^. Within the brain and VNC, only a single pair of neurons expressing ITP is located in the LN_d_ region (Fig. 3H). Analysis of the Fly SCope dataset suggests the presence of a restricted number of neurons that are triply positive for *ITP*, *NPF*, and *Clk* expression within the brain, but not in the VNC (Fig. S3K-N). Knockdown of *cyc*, induced apoptosis, hyperexcitation, or inhibition of synaptic transmission in ITP-LN_d_ and 5^th^ sLN_v_ neurons, consistently disrupted SMD behavior (Fig. S3E-I). However, inhibiting the neuronal activity of ITP-LN_d_ and 5^th^ sLN_v_ neurons via voltage-gated potassium channel, KCNJ2 resulted in developmental lethality, indicating that their neuronal function is essential for development as well (Fig. S3G). The feminization of ITP-LN_d_ and 5th sLN_v_ neurons through the expression of *UAS-tra^F^*had no effect on interval timing behavior (Fig. S3J), demonstrating that the sexual dimorphism of these LN_d_ neurons does not play a role in regulating interval timing. Notably, knockdown of *ITP* within ITP-LN_d_ neurons did not affect SMD behavior (Fig. 3G), suggesting that the role of ITP in ITP-LN_d_ neurons is distinct from its function in interval timing. These findings implicate a specialized subset of ITP-LN_d_ and 5^th^ sLN_v_ neurons as a critical node in the circuitry governing interval timing, while also highlighting the diverse functions of ITP in neuronal physiology.

### The glutamatergic output circuits originating from ITP-LN_d_ neurons are instrumental in the generation of interval timing

Our findings reveal that the expression of flippase in NPF-positive neurons with a UAS-stop cassette can confine the expression of *GAL4^R54D11^* to four specific neurons in the brain, including the ITP-LN_d_ and 5^th^-sLN_v_ neurons (Fig. 4A). Inhibiting the activity of these neurons disrupts SMD behavior without causing developmental lethality (Fig. 4B), indicating that the KCNJ2-mediated developmental lethality observed with *GAL4^R54D11^*is likely due to its expression in the VNC (Fig. S3G and Fig. 4A). Furthermore, hyperexcitation induced by NaChBac, inhibition of synaptic transmission by TNT, or the knockdown of *Clk* via *Clk-RNAi* in these four brain neurons all resulted in the disruption of SMD behavior (Fig. 4C-E), emphasizing the critical role of CLK function and neuronal activity in the ITP-LN_d_ neurons in the brain for the generation of interval timing.

**Figure 4.**
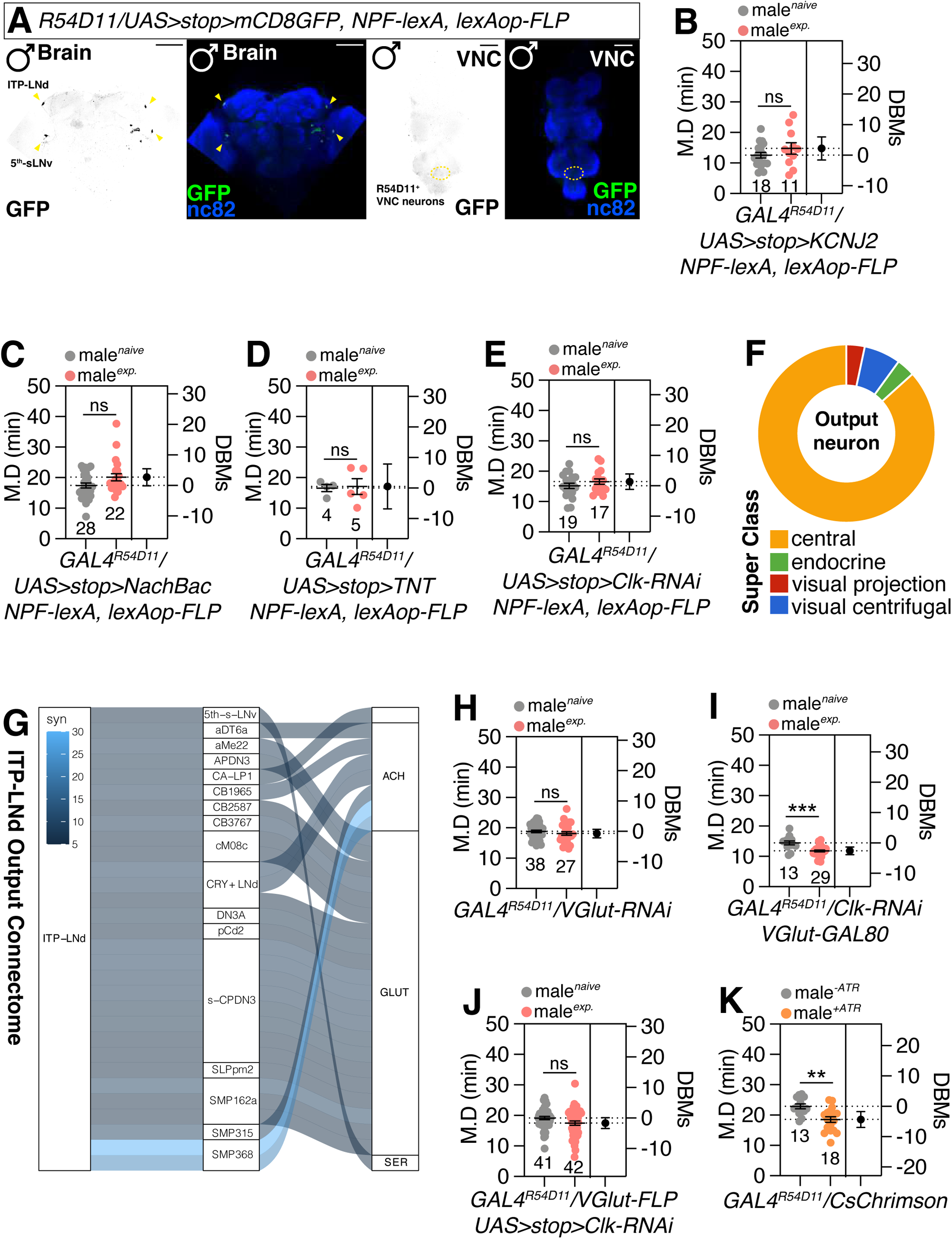
Normal glutamatergic output from the ITP-LN_d_ is necessary for interval timing. (A) Male flies expressing the *UAS(FRT.stop)mCD8GFP; GAL4^R54D11^* together with *lexAop-FLP; NPF-lexA* were immunostained with anti-GFP (green) and nc82 (magenta) antibodies. Yellow arrows indicate ITP-LN_d_ and 5^th^ sLN_v_ soma. Yellow dash circles indicate *GAL4^R54D11^*-labeled VNC neurons. Scale bars represent 50 μm. (B-E) SMD assays for the GAL4 system labeled only ITP-LN_d_ and 5^th^ sLN_v_ mediated (B) electrical silencing via *(FRT.stop)KCNJ2*, (C) electrical activating via *(FRT.stop)NaChBac*, (D) inactivation of synaptic transmission via *(FRT.stop)TNT*, and (E) knockdown of *Clk* via *(FRT.stop)Clk-RNAi*. (F-G) Connectome information of ITP-LN_d_ output neurons. (F) displays a donut diagram demonstrating the type of output neurons. (G) displays a Sankey histogram depicting the synaptic connections between ITP-LN_d_ and output neurons, the neurotransmitters that the output neurons use are displayed on the right. The color indicates the number of synapses connected. The data is from https://codex.flywire.ai/. The blank space in the upper right corner represents the failure to confirm the types of neurotransmitters used by these neurons. (H) SMD assays for *GAL4^R54D11^* driver mediated knockdown of vesicular glutamate transporter via *VGlut-RNAi*. (I) SMD assay for *GAL4^R54D11^*driver mediated knockdown of *Clk* in VGlut-negative neurons via *Clk-RNAi; VGlut-GAL80*. (J) SMD assay for *GAL4^R54D11^* driver mediated knockdown of *Clk* in glutamatergic neurons via *Clk-RNAi; VGlut^FLP^*. (K) Optogenetic experiment for *GAL4^R54D11^* driver mediated electrical activation via *UAS-CsChrimson*. Two groups of flies fed all-trans retinaldehyde (ATR) and unfed were irradiated overnight with red light to simulate cellular activation during SMD behavioral generation. See the “Methods” for a detailed description of the optogenetic experiment in this study.

Utilizing the fly SCope RNA seq dataset and the FlyWire connectome dataset platform, we inferred the expression patterns of enzymes responsible for neurotransmitter synthesis (Fig. S4A-B) and identified the output neurons from the ITP-LN ^34,41–48^. The FlyWire connectome data suggests that ITP-LN neurons project to central brain circuits (Fig. 4F) and slightly more to the contralateral brain (Fig. S4C). Although the Fly SCope data analysis was inconclusive (Fig. S4A-B), the FlyWire connectome dataset analysis clearly demonstrated that the output circuits from the ITP-LN_d_ neurons consist of glutamatergic and possibly cholinergic co-transmission (Fig. 4G). Given that the FlyWire connectome data predict the 5^th^ sLN_v_ to be serotonergic neurons (Fig. 4G) and that ITP-LN_d_ neurons have been confirmed to be non-cholinergic LN_d_ neurons^49^, we conclude that the single pair of ITP-LN_d_ neurons are glutamatergic.

The knockdown of *VGlut* specifically in these neurons disrupts SMD behavior, and the expression of *Clk-RNAi* in the non-glutamatergic subset of GAL4^R54D11^ neurons rescues the disrupted SMD behavior (Fig. 4H-I). Moreover, knockdown of *Clk* only in GAL4^R54D11^-positive and glutamatergic neurons is sufficient to disrupt SMD behavior (Fig. 4J), indicating the crucial role of Clk expression in a single pair of ITP-LN_d_ neurons. Optogenetic activation of GAL4^R54D11^ neurons can induce a reduction in mating duration without prior sexual experience (Fig. 4K), suggesting that artificial neuronal activation of these neurons can mimic the sexual experience-mediated internal states of pacemaker circuits.

### Variants of the CLK protein that undergo alternative splicing may dissociate circadian rhythms from interval timing within ITP-LN_d_ and 5^th^-sLN_v_ neurons

We have previously demonstrated that rival-induced prolonged mating duration (LMD), a distinct form of interval timing behavior in male flies, persists even in arrhythmic conditions such as continuous light exposure for five days^10^ (Fig. 5A). Similarly, SMD behavior remains intact under arrhythmic conditions (Fig. 5B). In the same condition, the rhythmic activity and sleep of flies was completely disorganized (Fig. S5A-B), indicating that interval timing behaviors are independent of the circadian rhythm. We previously conducted experiments at various times throughout the day with *Canton-S* flies and observed normal SMD behavior^11^, which is consistent with the reported results.

**Figure 5.**
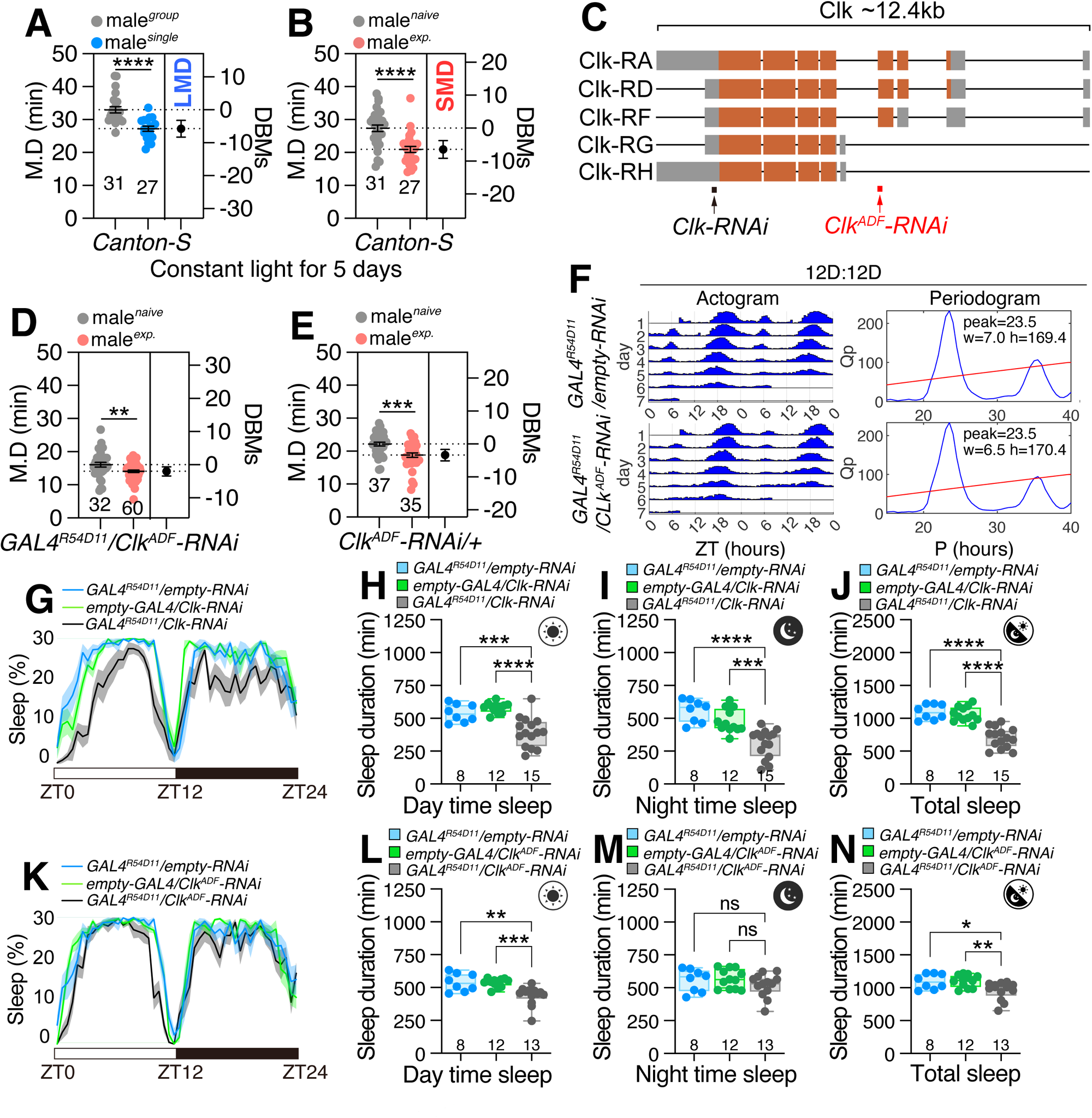
Different CLK protein isoforms in ITP-LN_d_ govern circadian rhythm and interval timing, respectively. (A-B) LMD and SMD assays for *Canton-S* males under 5 days of constant light. (C) Structures of the different *Clk* transcripts. Grey squares represent the exons, brown squares represent CDS, and arrows indicate short hairpin RNAs (shRNAs) for gene silencing. (D) SMD assay for *GAL4^R54D11^* driver mediated knockdown of *Clk-RA*, *RD*, and *RF* via *Clk^ADF^-RNAi*. (E) Genetic control of the SMD assay, which only has *Clk^ADF^-RNAi* without a GAL4 driver. (F) Actogram and Periodogram of single male flies for (top) genetic control (*GAL4^R54D11^* with *empty-RNAi*) and knockdown of *Clk-RA*, *RD*, and *RF* via *Clk^ADF^-RNAi* by *GAL4^R54D11^*(bottom). See the “Methods” for a detailed description of the single circadian rhythm experiment in this study. (G-J) Sleep profiles of single flies for *GAL4^R54D11^* mediated knockdown of *Clk* via *Clk-RNAi* and the quantification of sleep duration. The data were derived from the average sleep duration over a 3-5 day period under a 12-hour light:12-hour dark cycle. See the “Methods” for a detailed description of the single activity and sleep experiment in this study. (K-N) Sleep profiles of single flies for *GAL4^R54D11^* mediated knockdown of *Clk-RA*, *RD*, and *RF* via *Clk^ADF^-RNAi* and the quantification of sleep duration.

The influence of temperature and feeding on the alternative splicing of clock gene products has been explored^50–52^, but the role of CLK mRNA variants in regulating various timing behaviors has not been previously reported. The *Clk* gene produces five distinct mRNA transcripts, *Clk-RA*, *RD*, *RF*, *RG*, and *RH*, with *Clk-RG* and *RH* lacking the first three exons and thus unable to bind DNA. Our study revealed that *Clk^ADF^-RNAi* specifically targets *Clk-RA*, *RB*, and *RF* transcripts, which contain DNA-binding motifs. Surprisingly, the expression of *Clk^ADF^-RNAi* did not disrupt SMD behavior, or the circadian rhythm compared to genetic controls (Fig. 5D-F and Fig. S5C). This suggests that CLK proteins with DNA-binding motifs are not essential for generating interval timing behaviors in ITP-LN_d_ neurons.

We next investigated whether CLK expression in ITP-LN_d_ neurons is linked to sleep behaviors. Knockdown of *Clk* in ITP-LN_d_ neurons partially reduced the duration of sleep during both day and night (Fig. 5G-J). Remarkably, when we selectively knocked down *Clk* transcripts with DNA-binding motifs in ITP-LN_d_ neurons (Fig. 5C), daytime but not nighttime sleep duration decreased (Fig. 5K-N). This observation clearly indicates that different variants of CLK proteins have differential effects on sleep behavior. We speculate that CLK proteins lacking the DNA-binding motif may play a specific role in regulating sleep and interval timing, suggesting a novel molecular mechanism for the regulation of these behaviors.

### The CLK/CYC heterodimer, a transcription factor complex, is specifically linked to interval timing and sleep behaviors

Although circadian clock genes are primarily known for their role in regulating circadian rhythms, they also perform various other functions^53–55^. Sleep timing is influenced by the circadian clock, yet the two processes can be dissociated; mutations in core clock genes often lead to fragmented sleep patterns but do not necessarily affect overall sleep duration^28^. The *cyc^01^* and *Clk^Jrk^* mutants displayed distinct sleep pattern abnormalities and life span changes, with the *cyc^01^*mutation exhibiting sexually dimorphic effects on sleep compensation^56^. The CLK/CYC heterodimer transcription factors play a pivotal role in the regulation of numerous clock gene transcripts. However, the specific functions of these target genes in various timing behaviors are not yet fully elucidated (Fig. 6A)^23,57^.

**Figure 6.**
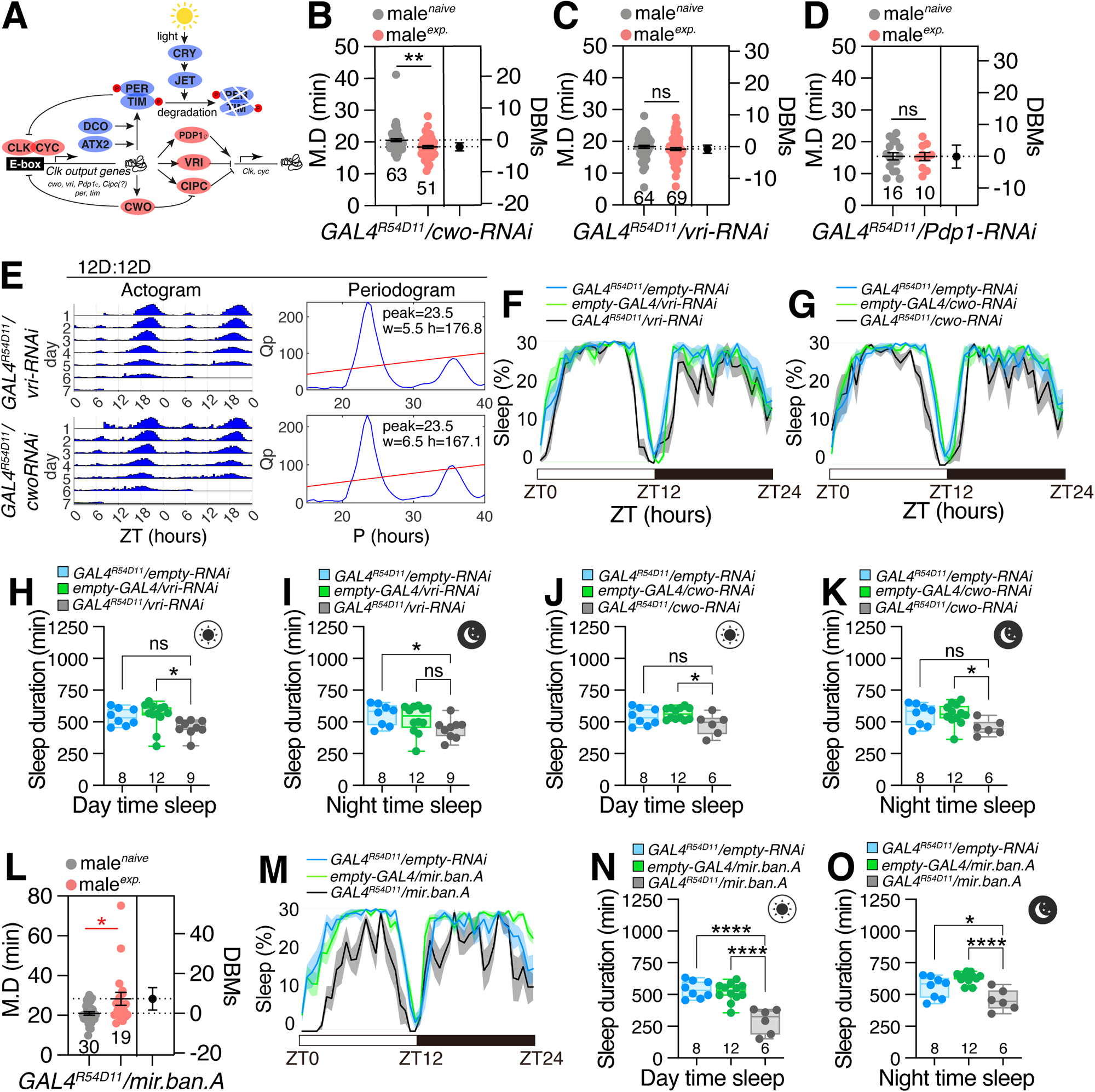
Different circadian genes produce distinct effects on interval timing and sleep behavior. (A) Diagram of the molecular clock. Red ovals represent CLK/CYC-associated proteins, and blue ovals represent PER/TIM-associated proteins. (B-D) SMD assay for *GAL4^R54D11^* driver mediated knockdown of CLK/CYC-associated genes via (B) *cwo-RNAi*, (C) *vri-RNAi*, and (D) *Pdp1-RNAi*. (E) Actogram and Periodogram of single male flies for *GAL4^R54D11^* driver mediated knockdown of *vri* or *cwo* via *cwo-RNAi* (top) or *vri-RNAi* (bottom). (F-G) Sleep profiles of flies for *GAL4^R54D11^* mediated knockdown of *vri* or *cwo* via (F) *vri-RNAi* or (G) *cwo-RNAi*. (H-K) Quantification of sleep duration for *GAL4^R54D11^* mediated knockdown of *vri* or *cwo* via (H-I) *vri-RNAi* or (J-K) *cwo-RNAi*. (G) SMD assay for *GAL4^R54D11^* driver mediated overexpression of microRNA *bantam* via *UAS-mir.ban.A*. (M-O) Sleep profiles and quantification of flies for *GAL4^R54D11^* mediated overexpression of microRNA *bantam* via *UAS-mir.ban.A*.

Our findings suggest that the CLK/CYC heterodimer lacking a DNA-binding motif is a critical regulator of interval timing and sleep, but not necessarily of circadian rhythm. Consequently, we selected several essential clock genes known to modulate circadian rhythm and reassessed their roles in interval timing and sleep within ITP-LN_d_ neurons. Knockdown of *cwo* (*clockwork orange*) and *Cipc* (*Clock interacting protein circadian*) had no impact on SMD behavior, while *vri* (*vrille*) and *Pdp1* (*PAR-domain protein 1*) knockdown disrupted SMD (Fig. 6B-D and Fig. S6A). Notably, PER/TIM-related clock genes, such as *dco* (*discs overgrown*), *jet* (*jetlag*), *Atx2* (*Ataxin-2*), and *cry*, did not affect SMD behavior (Fig. S6B-E). The expression levels of these genes did not correlate with their influence on SMD (Fig. S6F).

To further investigate the impact of CLK/CYC function on circadian rhythm and sleep within ITP-LN_d_ neurons, we selected *cwo* and *vri* as representative genes. Both genes are closely associated with CLK/CYC function, but only *vri* is involved in interval timing. Knockdown of either genes did not disrupt circadian rhythm (Fig. 6E and Fig. S6G). Knockdown of *vri* in ITP-LN_d_ neurons resulted in a decrease in daytime and nighttime sleep duration (Fig. 6F and Fig. 6H-I), while *cwo* knockdown reduced the duration of sleep similarly (Fig. 6J and Fig. 6J-K). These results indicate that CLK-VRI interaction is crucial for modulating interval timing and sleep, but not for circadian rhythm regulation, within ITP-LN_d_ neurons. Overexpression of miRNA *bantam*, which binds to the 3’ UTR of Clk and modulates its translation^58^, reversed SMD behavior, indicating the critical role of CLK protein regulation in generating interval timing (Fig. 6L). Concurrently, the overexpression of *mir.ban* in ITP-LN_d_ resulted in significant alterations in sleep behavior (Fig. 6M). Specifically, this was manifested as a decrease in sleep during the daytime and an increase during the nighttime (Fig. 6N-O). These findings suggest that *mir.ban* also plays a pivotal role in the regulation of sleep behavior.

### The SIFa-SIFaR peptidergic signaling pathway is specifically involved in modulating interval timing distinct from its influence on circadian rhythm or sleep behavior

Circadian clock cells are influenced by environmental factors such as temperature, light, and feeding^53,55,59–61^. To elucidate the input signals that specifically regulate SMD behavior in ITP-LN_d_ neurons, we analyzed the fly SCope dataset, which revealed that SIFaR is highly expressed in these neurons (Fig. 7A). Neuropeptide SIFa, expressed in four neurons in the *pars intercerebralis* (PI), plays a pivotal role in modulating interval timing behaviors^12^. Knockdown of *SIFaR* in ITP-LN_d_ neurons, as well as knockdown of *Clk* in SIFaR-positive cells, disrupted SMD behavior (Fig. 7B-C), but did not affect circadian rhythm (Fig. S7A). Concomitantly, the absence of SIFaR led to a reduction of both daytime and nighttime sleep duration (Fig. S7B-E). BAcTrace data demonstrated that GAL4^R54D11^ neurons receive strong input signals from PI-located SIFa neurons^62^ (Fig. 7D). These findings suggest that SIFa signals are crucial for interval timing and sleep regulation in ITP-LN_d_ neurons, but not for circadian rhythm.

**Figure 7.**
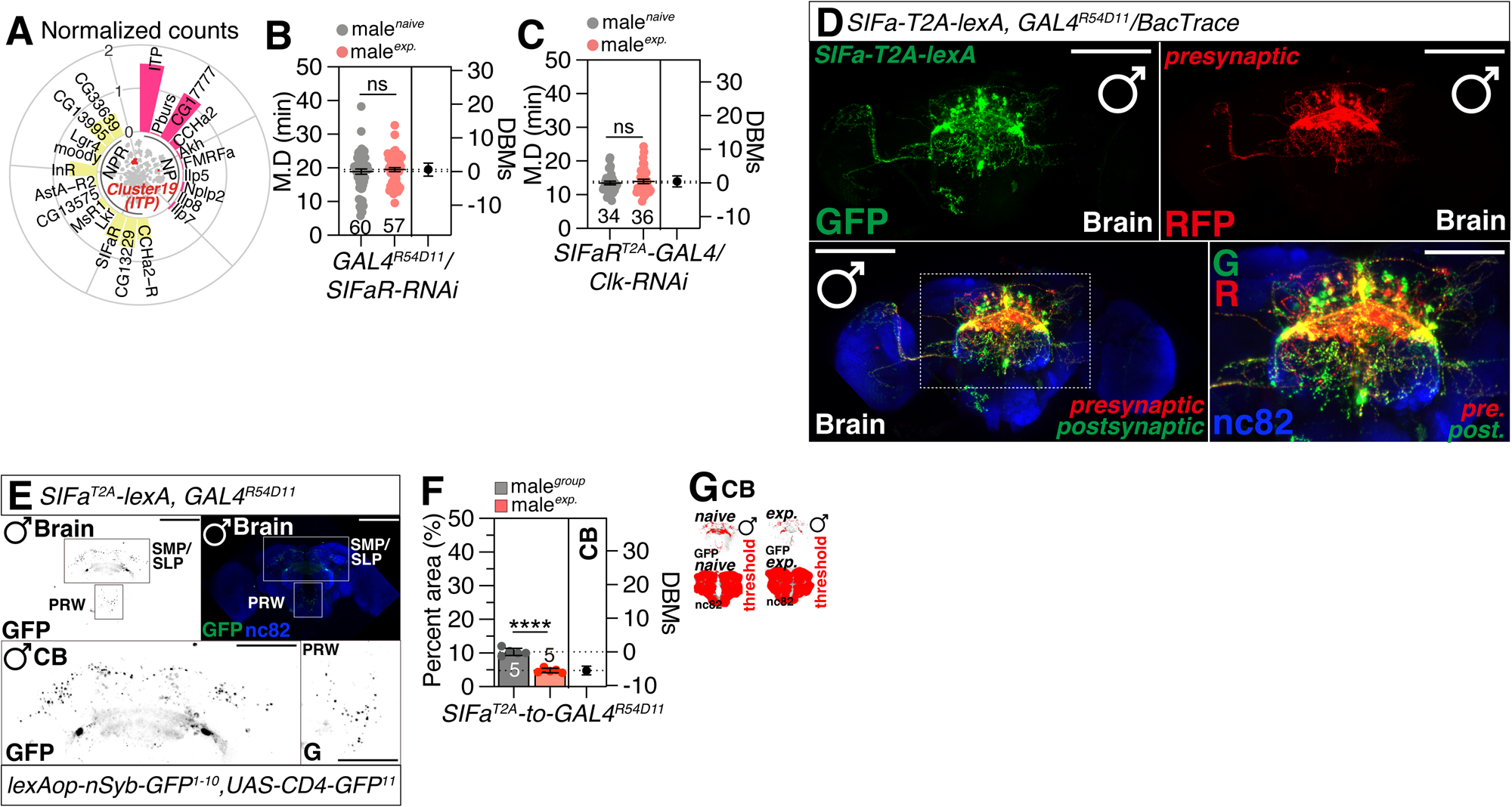
The SIFa-SIFaR peptidergic signaling pathway regulates the formation of interval timing behavior. (A) Expression of neuropeptide (NP) and neuropeptide receptor (NPR) in ITP-LN cells. The average UMI count was calculated for each gene in this population, followed by log10 transformation for data normalization. To focus on cells expressing these genes, only genes exhibiting increased expression of NP and NPR were included. (B) SMD assay for *SIFaR^T2A^-GAL4* driver mediated knockdown of *Clk* in SIFaR-positive neurons via *Clk-RNAi*. (C) SMD assay for *SIFaR^T2A^-GAL4* driver mediated knockdown of *Clk* in SIFaR-positive neurons via *Clk-RNAi*. (D) Male flies expressing *SIFa^T2A^-lexA; GAL4^R54D11^* together with a retro-grade and transsynaptic labeling system, *BAcTrace*, were immunostained with anti-GFP (green), anti-DsRed (red), and nc82 (magenta) antibodies. Left: expression pattern of *SIFa^T2A^-lexA*, middle: BAcTrace transsynaptic-labeled presynaptic neurons of GAL4^R54D11^ neurons; right: SIFa neurons overlap with the BAcTrace-labeled presynaptic neurons of GAL4^R54D11^ neurons. Scale bars represent 50 μm. (E) GRASP assay for *SIFa^2A-lexA^* and *GAL4^R54D11^* together with *lexAop-nsyb-spGFP^1-10^, UAS-CD4-spGFP^11^* in the SLP and SMP regions of naïve male flies. Scale bars represent 50 μm. Brains of male fly were immunostained with anti-GFP (green) and anti-nc82 (blue) antibodies. The left and bottom panels are presented as a gray scale to clearly show the synapses connection between *SIFa^2A-lexA^* and *GAL4^R54D11^*. Boxes indicate the magnified regions of interest presented in the bottom panels. (F-G) Quantification of relative value for synaptic areas that are formed between *SIFa^2A-lexA^* and *GAL4^R54D11^* in (E, S8D) between naïve and single male flies. The same quantification was performed for the relative synaptic area in these brain regions between naïve and experienced male flies. The synaptic interactions were visualized utilizing the GRASP system in naïve and experienced male flies. SMP, superior medial protocerebrum; SLP, superior lateral protocerebrum; CB, central brain; PRW, prow. (G) The small panels are presented as a red scale to show the GFP fluorescence marked by the threshold function of ImageJ. See the “Methods” for a detailed description of the fluorescence intensity analysis used in this study.

Furthermore, we observed that ITP-LN_d_ neurons form strong synapses with SIFa neurons in the superior medial protocerebrum (SMP) and superior lateral protocerebrum (SLP) regions of the brain (Fig. 7E). The SLP region is known to be involved in upstream processing of mushroom body (MB) learning centers and is where long-range taste projection neurons project^63^. Although sexual experiences did not significantly alter the accumulated calcium levels in ITP-LN_d_ neurons (Fig. S8A-B) and activation of SIFa neurons also did not lead to elevated calcium levels in ITP-LN_d_ (Fig. S8C), the synapses between SIFa and ITP-LN_d_ neurons were significantly reduced (Fig. 7F-G and Fig. S8D). These data indicate that sexual experiences reduce the inputs to ITP-LN_d_ neurons from SIFa peptidergic signals without dramatic calcium flux change.

Using the FlyWire dataset, we analyzed the input circuits to ITP-LN_d_ neurons and identified that cholinergic, glutamatergic, and GABAergic inputs constitute the main neurotransmitters from input neurons, with half coming from visual and half from central class neurons (Fig. S9A-B). Interestingly, the inputs to ITP-LN_d_ neurons predominantly originate from the left side of the brain, potentially representing the involvement of the asymmetric body (AB) associated with this circuit (Fig. S9C)^64–68^.

The synaptic connectivity of ITP-LN_d_ neurons has previously been shown to be distinct within the clock network, exhibiting fewer intra-network synapses compared to other groups of clock neurons ^69^. Previous single-cell RNA sequencing data analysis of 150 clock neurons identified that neural connectivity molecules are key in defining the diverse functions of heterogeneous clock cells^70^. Among these molecules, voltage-gated potassium channel Shaker (Sh) and its beta-subunit Hyperkinetic (Hk) were found to be highly enriched in ITP-LN_d_ neurons. Knockdown of these channels in ITP-LN_d_ neurons reversed SMD behavior, indicating their critical role in maintaining ITP-LN_d_ as pacemaker neurons for interval timing (Fig. S9D-E). Interestingly, the reversed phenotype of interval timing associated with Sh knockdown is linked to a significant shortening of sleep duration, while Hk knockdown shows no effect on sleep. (Fig. S9F-M), suggesting that these channels play critical roles in ITP-LN_d_ neurons, specifically modulating interval timing and sleep independently of circadian rhythms.

Knockdown of the Ih channel, which encodes a low-threshold, voltage-gated ion channel in ITP-LN_d_, also disrupted SMD behavior (Fig. S9N), suggesting that voltage-gated channels are important regulators of pacemaker function for interval timing. In contrast, knockdown of serotonin receptor 5-HT1A, octopamine receptor in MB (Oamb), or octopamine b3 receptor (Octb3R), which are known to be enriched in ITP-LN_d_ neurons^70^, did not affect SMD behavior (Fig. S9O-Q), indicating that 5-HT and octopamine (OA) transmission are not essential for ITP-LN_d_’s role in interval timing. These data collectively underscore the pivotal role of voltage-gated channels in maintaining the pacemaker function of CLK-expressing ITP-LN_d_ neurons for interval timing.

## Discussion

In this study, we investigated the role of the CLK/CYC heterodimer and associated factors within a specific pair of LN_d_ neurons in modulating sexual experience-dependent interval timing (SMD) behavior. We found that mutations in *Clk* and *cyc*, but not *per* and *tim*, disrupt SMD behavior, indicating a unique role for CLK/CYC in interval timing. Neuronal CLK/CYC expression is necessary and sufficient for SMD, while glial expression is not (Fig. 1). Specific CLK/CYC expression in NPF^+^ and cry^+^ neurons in the brain, but not in PDF neurons, is essential for SMD (Fig. 2). Detailed analysis identifies CLK expression in ITP-LN_d_ and 5^th^ sLN_v_ neurons as pivotal for SMD. Knockdown of CLK in these neurons impairs SMD, as does inhibiting their activity (Fig. 3). ITP-LN_d_ neurons are glutamatergic, with output circuits to central brain. Inhibiting glutamatergic transmission or knocking down CLK in these neurons disrupts SMD. Furthermore, optogenetic activation of these neurons can induce SMD without prior experience (Fig. 4). We also explored the molecular mechanisms, finding that CLK variants lacking DNA binding motifs dissociate circadian rhythms from interval timing and sleep behaviors in ITP-LN_d_ neurons (Fig. 5). CLK-VRI interaction is crucial for modulating interval timing and sleep, but not circadian rhythm, within ITP-LN_d_ neurons (Fig. 6). Finally, we demonstrated that the SIFa-SIFaR peptidergic signaling pathway is specifically involved in modulating interval timing distinct from its influence on circadian rhythm (Fig. 7). Our study provides a comprehensive understanding of the role of CLK/CYC heterodimer and associated factors in modulating interval timing behavior in *Drosophila*, shedding light on the neural circuits and molecular mechanisms underlying this complex behavior (Fig. S9R).

Despite the central role of circadian clock genes in regulating circadian rhythms, these genes are also involved in various non-circadian functions. Accumulating evidence suggests that clock genes exhibit pleiotropic effects in regulating diverse physiological processes^53,55,59,60^. In addition to their well-established roles in sleep homeostasis^28,53,59,71–73^, clock genes have been implicated in the modulation of addiction, psychiatric disorders, and male fertility^53,74,75^. Recent studies have expanded our understanding of CLK’s function, revealing its role in tissue-specific gene regulation and metabolic processes^76^, with Gart being a notable example that is crucial for maintaining feeding rhythms and food intake^73^. Furthermore, distinct clock neurons have been shown to mediate the circadian rhythm of male sex drive^77^. The surprising diversity of circadian clock cells, akin to their functional heterogeneity, suggests that individual clock neurons possess distinct functions beyond circadian rhythm regulation^78^. In conclusion, the diverse non-circadian functions of clock genes underscore their intricate regulatory roles across various physiological processes. The discovery of specific non-circadian functions of clock genes, such as the role of CLK in modulating interval timing and sleep regulation in ITP-LN_d_ neurons, offers valuable insights into the multifaceted functions of these genes. These findings underscore the complexity and versatility of clock genes in orchestrating biological rhythms and behaviors beyond the circadian clock.

In our study, we uncovered that various *Clk* transcripts exhibit distinct timing functions. Our data indicate that *Clk-RA*, *RD*, and *RF* are essential for circadian rhythmicity and sleep, while *Clk-RG* and *RH* are critical for interval timing and sleep regulation. These findings suggest that *Clk* transcripts regulate a spectrum of timing behaviors through their expression in specific clock cells. Moreover, we identified voltage-gated potassium channels Sh and Hk as pivotal for maintaining the pacemaker function of ITP-LN_d_ neurons in regulating interval timing. Elucidating the relationship between Clk-RG/RH function and Sh/Hk in ITP-LN_d_ neurons may provide insights into the molecular mechanisms governing pacemaker regulation in the fly brain for interval timing behavior (Fig. S9R).

The traditional perspective on the rhythmic regulation of CLK/CYC dimer-mediated transcriptional control does not account for the PER/TIM-independent mechanisms or the specific dependence of SMD behavior on alternative splicing forms of *Clk*, which are crucial for behavioral timekeeping within a 5-20-minute range. Our data suggest that a C-terminally deleted CLK protein can exhibit circadian-independent functions, particularly in relation to male interval timing associated with mating investment behavior. Recent studies have reported that alternative splicing of the *Clk* transcript mediates the circadian clock’s response to temperature changes^79^. Additionally, a non-circadian role for *cyc* has been identified, which regulates the development of clock neurons^80^. There has long been speculation regarding the non-circadian regulation and function of clock genes in controlling oogenesis in female *Drosophila*^81^ and in influencing non-circadian aspects of mating behavior^55^. Therefore, we propose that CLK/CYC dimers composed of different forms of CLK protein may serve diverse functions beyond circadian rhythm regulation, including metabolic control, feeding behavior, mating behavior, and interval timing. Further investigation into the various forms of CLK proteins and their dimerization with CYC will shed light on how these non-circadian functions have evolved alongside the circadian roles of CLK protein.

## Supporting information

Table 1. Screen of 150 clock neurons.

Graphical Abstract

## Acknowledgments

We thank Drs. Yuh Nung Jan and Lily Yeh Jan (UCSF, USA) for helpful comments and support on this paper. We are very appreciative to the colleagues who supplied us with several fly strains: Dr. Alex C. Keene (Texas A&M University), Dr. Justin Blau (New York University), Dr. Amita Sehgal (University of Pennsylvania), Dr. Ravi Allada (University of Michigan), Dr. Lihua Jin (Northeast Forestry University), Dr. Zongzhao Zhai (Hunan Normal University), Dr. Wei Zhang (Tsinghua University), Dr. Donggen Luo (Peking University), Dr. Fang Guo (Zhejiang University), Dr. Yufeng Pan and Dr. Junhai Han (Southeast University) and Drs. Young-Joon Kim and Sung-Eun Yoon (Korea Drosophila Resource Center, KDRC). This research was supported a University of Ottawa Startup grant 602496 to WJK, Startup funds from HIT Center for Life Science to WJK, a University of Ottawa Interdisciplinary Research Group Funding Opportunity (IRGFO stream 1 and 2) grants 148101 and 148747 to WJK, a Natural Sciences and Engineering Research Council of Canada (NSERC) Discovery grant (reference: 211406) to WJK, a University of Ottawa Brain and Mind Research Institute/Center for Neural Dynamics Open call project grant 150950 to WJK, a Mitacs Globalink Research Internship Program grant 17268 to WJK. This research was also supported by the Brain Pool Program of the National Research Foundation in Korea grant ZYM5041911 to WJK, Burroughs Wellcome Fund Collaborative Research Travel Grants (reference: 1017486) to WJK and a NVIDIA Academic Hardware Grant Program to WJK. The funders had no role in study design, data collection and analysis, decision to publish, or preparation of the manuscript. HM received salary from the ‘Startup funds from HIT Center for Life Science to WJK’.

## Author Contributions

**Conceptualization:** Woo Jae Kim.

**Data curation:** Hongyu Miao, Zekun Wu, Yanan Wei, Woo Jae Kim.

**Formal analysis:** Hongyu Miao, Zekun Wu, Woo Jae Kim.

**Funding acquisition:** Woo Jae Kim.

**Investigation:** Woo Jae Kim.

**Methodology:** Woo Jae Kim.

**Project administration:** Woo Jae Kim.

**Resources:** Woo Jae Kim.

**Supervision:** Woo Jae Kim.

**Validation:** Hongyu Miao, Woo Jae Kim.

**Visualization:** Hongyu Miao, Zekun Wu, Woo Jae Kim.

**Writing – original draft:** Woo Jae Kim.

**Writing – review & editing:** Hongyu Miao, Woo Jae Kim.

## Declaration of Generative AI and AI-assisted Technologies in the Writing Process

During the creation of this work, the author(s) utilized QuillBot to rephrase English sentences, verify English grammar, and detect plagiarism, as none of the authors of this paper are native English speakers. After using this tool/service, the author(s) reviewed and edited the content as needed and take(s) full responsibility for the content of the publication.

**Table 1.** Screen of 150 clock neurons.

## Resource Availability

### Lead contact

Further information and requests for resources and reagents should be directed to and will be fulfilled by the lead contact, Woo Jae Kim.

## Data and code availability

- All data reported in this paper will be shared by the lead contact upon request.
- This paper does not report original code. The URL of the codes used in this paper are listed in the key resources table.
- Any additional information required to reanalyze the data in this paper is available from the lead contact upon request.

## Methods

### Fly stocks and husbandry

*Drosophila melanogaster* were raised on cornmeal-yeast medium at similar densities to yield adults with similar body sizes. Flies were kept in 12 h light: 12 h dark cycles (LD) at 25L (ZT 0 is the beginning of the light phase, ZT12 beginning of the dark phase) except for some experimental manipulation (experiments with the flies carrying tub-GAL80^ts^). Wild-type flies were *Canton-S*. To reduce the variation from genetic background, all flies were backcrossed for at least 3 generations to CS strain. All mutants and transgenic lines used here have been described previously.

The following lines were obtained from Dr. Alex C. Keene (Texas A&M University) and Dr. Justin Blau (New York University): *cry^03^*, *cry-GAL80*, *tim-GAL4* (BDSC7126).

The following lines were obtained from Dr. Amita Sehgal (University of Pennsylvania): *qvr^1^*.

The following lines were obtained from Dr. Lihua Jin (Northeast Forestry University): *Myo1A-GAL4*, *esg-GAL4*, *Hml-GAL4* (BDSC30139).

The following lines were obtained from Dr. Zongzhao Zhai (Hunan Normal University): *uro-GAL4* (BDSC91415).

The following lines were obtained from Dr. Wei Zhang (Tsinghua University): *tsh-GAL80* (BDSC605556), *lexAop-FLP* (BDSC55819), *VGlut-GAL80* (BDSC58448).

The following lines were obtained from Dr. Ravi Allada (University of Michigan): *UAS-cyc*.

The following lines were obtained from Dr. Donggen Luo (Peking University) and Dr. Junhai Han (Southeast University): *Clk4.1M-GAL4* (BDSC31316), *GAL4^R54D11^*(BDSC41279), *Mai179-GAL4*.

The following lines were obtained from Dr. Yufeng Pan (Southeast University): *UAS-jGCaMP7s* (BDSC79032), *empty-RNAi* (BDSC36304).

The following lines were obtained from Bloomington Stock Center: *Canton-S* (64349), *Df(1)Exel6234* (7708), *per^01^* (80928), *tim^01^* (80930), *Clk^Jrk^*(80927), *cyc^01^* (80929), *elav^c155^* (458), *Clk-RNAi^HMJ02224^* (42566), *cyc-RNAi^HMJ02219^* (42563), *repo-GAL4* (7415), *grh-GAL4* (65637), *Mhc-GAL4* (55133), *Clk-RNAi^JF01453^*(31660), *Clk-RNAi^JF01454^* (31661), *cyc-RNAi^JF03333^*(29400), *cyc-RNAi^JF02185^* (31897), *cyc-RNAi^GL00387^*(35461), *tub(FRT.GAL80)* (38881), *otdFLP* (600309), *cry-GAL4; Pdf-GAL80* (80940), *cry-GAL4* (24514), *NPF-GAL4* (25681), *NPF-GAL4* (25682), *tub-GAL80^ts^* (7108), *UAS-CD4tdGFP* (35839), *UAS-RedStinger* (8546), *ITP-RNAi* (25799), *UAS-hid* (65403), *UAS-KCNJ2* (6595), *UAS-NaChBac* (9469), *UAS-TNT* (28838), *UAS-traF* (4590), *UAS>stop>KCNJ2* (67686), *VGlut-RNAi* (27538), *UAS-CsChrimson* (55136), *cwo-RNAi* (26318), *vri-RNAi* (40862), *Pdp1-RNAi* (26212), *BacTrace* (90826), *SIFaR-RNAi* (34947), *lexAop-nSyb-spGFP^1-10^, UAS-CD4-spGFP^11^* (64315), *Cipc-RNAi* (28774), *dco-RNAi* (27719), *jet-RNAi* (31058), *Atx2-RNAi* (36114), *cry-RNAi* (43217), *UAS-mir.ban.A* (60671), *lexAop-CD8GFP; UAS-mLexA-VP16-NFAT, lexAop-rCD2-GFP* (66542), *Sh-RNAi* (53347), *Hk-RNAi* (28330), *Oamb-RNAi* (31233), *UAS-mCD8RFP, LexAop-mCD8GFP, nSyb-MKII::nlsLexADBDo, UAS-p65AD::CaM* (61679), *empty-GAL4* (36303).

The following lines were obtained from Qidong Fungene Biotechnology: *ITP-RC^T2A^-GAL4* (FBA00286), *VGlut^FLP^* (FRE00001), *SIFa-lexA^T2A^*(FBF00116).

The following lines were obtained from Korea Drosophila Resource Center: *UAS>stop>mCD8GFP* (1119), *UAS>stop>NaChBac* (1183), *UAS>stop>TNT* (1191).

The following lines were obtained from Vienna Drosophila Resource Center: *Clk^ADF^-RNAi* (104507), *5-HT1A-RNAi* (106094).

The following lines were obtained from TsingHua Fly Center: *Ih-RNAi* (TH02084.N).

The following lines were obtained from NIG-FLY Center: *Oct*β*3R-RNAi* (31348R-4).

The CS background was selected as the experimental background due to its well-characterized and consistent LMD and SMD behaviors. To ensure that genetic variation did not confound our results, all GAL4, UAS, and RNAi lines employed in our assays were rigorously backcrossed into the CS strain, often exceeding ten generations of backcrossing. This approach was undertaken to isolate the effects of our genetic manipulations from those of genetic background. We assert that the extensive backcrossing to the CS background, in concert with the internal control in LMD and SMD, provides a stable platform for the accurate interpretation of the LMD and SMD phenotypes observed in our experiments. To reduce the variation from genetic background, all flies were backcrossed for at least 10 generations to *CS* strain. For the generation of outcrosses, all GAL4, UAS, and RNAi lines employed as the virgin female stock were backcrossed to the *CS* genetic background for a minimum of ten generations. Notably, the majority of these lines, which were utilized for LMD assays, have been maintained in a *CS* backcrossed state for long-term generations subsequent to the initial outcrossing process, exceeding ten backcrosses. Based on our experimental observations, the genetic background of primary significance is that of the X chromosome inherited from the female parent. Consequently, we consistently utilized these fully outcrossed females as virgins for the execution of experiments pertaining to LMD and SMD behaviors. Contrary to the influence on LMD behaviors, we have previously demonstrated that the genetic background exerts negligible influence on SMD behaviors, as reported in our prior publication^11^. The mutants and transgenic lines utilized in this study have been previously characterized, with the exception of the novel transgenic strains that we generated and describe herein.

## Mating duration assay

The mating duration assay in this study has been reported^9–11^. To enhance the efficiency of the mating duration assay, we utilized the *Df(1)Exel6234* (DF here after) genetic modified fly line in this study, which harbors a deletion of a specific genomic region that includes the sex peptide receptor (SPR)^82,83^. Previous studies have demonstrated that virgin females of this line exhibit increased receptivity to males^83^. We conducted a comparative analysis between the virgin females of this line and the CS virgin females and found that both groups induced SMD. Consequently, we have elected to employ virgin females from this modified line in all subsequent studies. For group reared (naïve) males, 40 males from the same strain were placed into a vial with food for 5 days. For single reared males, males of the same strain were collected individually and placed into vials with food for 5 days. For experienced males, 40 males from the same strain were placed into a vial with food for 4 days then 80 DF virgin females were introduced into vials for last 1 day before assay. 40 DF virgin females were collected from bottles and placed into a vial for 5 days. These females provide both sexually experienced partners and mating partners for mating duration assays. At the fifth day after eclosion, males of the appropriate strain and DF virgin females were mildly anaesthetized by CO_2_. After placing a single female in to the mating chamber, we inserted a transparent film then placed a single male to the other side of the film in each chamber. After allowing for 1 h of recovery in the mating chamber in 25L incubators, we removed the transparent film and recorded the mating activities. Only those males that succeeded to mate within 1 h were included for analyses. The initiation and completion of copulation were recorded to the nearest second, with a precision of ±10 seconds. The total mating duration for each pair was determined from the moment of successful genital apposition until the separation of the male and female *Drosophila*.

Genetic controls with *GAL4/+* or *UAS/+* lines were omitted from supplementary figures, as prior data confirm their consistent exhibition of normal LMD and SMD behaviors^9–11,14,16^. Hence, genetic controls for LMD and SMD behaviors were incorporated exclusively when assessing novel fly strains that had not previously been examined. In essence, internal controls were predominantly employed in the experiments, as LMD and SMD behaviors exhibit enhanced statistical significance when internally controlled. Within the LMD assay, both group and single conditions function reciprocally as internal controls. A significant distinction between the naïve and single conditions implies that the experimental manipulation does not affect LMD. Conversely, the lack of a significant discrepancy suggests that the manipulation does influence LMD. In the context of SMD experiments, the naïve condition (equivalent to the group condition in the LMD assay) and sexually experienced males act as mutual internal controls for one another. A statistically significant divergence between naïve and experienced males indicates that the experimental procedure does not alter SMD. Conversely, the absence of a statistically significant difference suggests that the manipulation does impact SMD. Hence, we incorporated supplementary genetic control experiments solely if they deemed indispensable for testing. All assays were performed from noon to 4 PM. We conducted blinded studies for every test.

## Generation of transgenic flies

To generate the *UAS>stop>Clk-RNAi* line, we selected HMJ02224 (BDSC#42566) as the template shRNA. The shRNA sequences were cloned directly with the following primers TTATCCCATATTCAGCCGCTAGCAGT-AGAGCTAGTTGTAGATCTCAA-TAG TTATATTCAAGC and AACTCCGATGTCTCGCCTGAATTCGC-AGAGCTAGTTGTAGATCTCAA-TAT GCTTGAATATAAC. The amplified DNA fragment was inserted into the pJFRC28-10XUAS-FRT-stop-FRT-RNAi vector. This vector, supplied by Qidong Fungene Biotechnology Co., Ltd. (http://www.fungene.tech/), is a derivative of the pJFRC28-10XUAS-IVS-GFP-p10 vector (available at https://www.addgene.org/36431). The insertion was achieved by digesting the fragment and the vector with EcoRI and NheI restriction enzymes to create compatible sticky ends. The genetic construct was inserted into the *attp5* site on chromosome II and *VK0005* site on chromosome III to generate transgenic flies using established techniques, a service conducted by Qidong Fungene Biotechnology Co., Ltd.

## Immunostaining

After 5 days of eclosion, the Drosophila brain was taken from adult flies and fixed in 4% formaldehyde at room temperature for 30 minutes. The sample was than washed three times (5 minutes each) in 1% PBT and then blocked in 5% normal goat serum for 30 minutes. Subsequently, the sample was incubated overnight at 4L with primary antibodies in 1% PBT, followed by the addition of fluorophore-conjugated secondary antibodies for one hour at room temperature. Finally, the brain was mounted on plates with an antifade mounting solution (Solarbio) for imaging purposes. Samples were imaged with Zeiss LSM880. Antibodies were used at the following dilutions: Chicken anti-GFP (1:500, Invitrogen), mouse anti-nc82 (1:50, DSHB), rabbit anti-DsRed (1:500, Rockland Immunochemicals), Alexa-488 donkey anti-chicken (1:200, Jackson ImmunoResearch), Alexa-555 goat anti-rabbit (1:200, Invitrogen), Alexa-647 goat anti-mouse (1:200, Jackson ImmunoResearch).

## Quantitative analysis of fluorescence intensity

To quantify the calcium level and synaptic intensity in microscopic images, we introduced ImageJ software^84^. We initially employed ImageJ’s ‘Measure’ feature to calculate average pixel intensity across the entire image or in user-specified sections, and the ‘Plot Profile’ feature to create intensity profiles across areas. To maximize precision, we converted color images to grayscale before analysis. Thresholding methods were also utilized to produce binary images that accurately outlined areas of interest, with pixel intensities of 255 (white) assigned to regions of interest and 0 (black) to the background. Intensity values from the binary image were then transferred to the corresponding locations in the original grayscale image to obtain precise intensity measurements for each object. The ‘Display Results’ feature provided comprehensive data for each object, including average intensity, size, and other relevant statistics. To normalize for fluorescence differences between ROIs, GFP fluorescence for GRASP was normalized to nc82. All specimens were imaged under identical conditions.

## Optogenetic experiment

A 100LmM stock solution of all-trans-retinal (ATR) powder (Sigma) was prepared by dissolving it in 100% alcohol. For optogenetic experiments, 250 μl of the stock solution was mixed with 30Lml of 5% sucrose and 1% agar medium to prepare food with a final concentration of 400 μM ATR. Flies aged between 3 and 5 days were transferred to ATR food for a minimum of 3 days before performing optogenetic experiments^85^. ATR-fed flies and unfed flies were housed in separate transparent tubes and exposed to a 20s red light: 40s no-light cycle treatment overnight before mating duration assay.

## GCaMP experiment

Fly anesthesia was induced using CO_2_. Dissecting the brain after feeding ATR for at least three days. AHL solution (108 mM NaCl, 5 mM KCl, 2LmM MgCl_2_, 2LmM CaCl_2_, 4LmM NaHCO_3_, 1LmM NaH_2_PO_4_, 5 mM HEPES, 10 mM Sucrose, 5LmM Trehalose, pH 7.5) was used for both dissection and imaging to maintain neuronal activity. We then used Zeiss LSM880 confocal microscope to record calcium signaling fluctuations in ITP-LN_d_, 5^th^ sLN_v_, and ocelli in parallel with activation of SIFa neurons using 5 seconds of red light. The brains were scanned at 1 Hz sampling rate with the max pinhole. Fiji was used to examine ROIs.

ΔF/F_0_L=L(F_t_L−LF_0_)/F_0_L×L100%. F_0_ was the averaged fluorescence of the baseline.

Because laser irradiation causes the fluorescence signal to diminish over time, we used the R package feasts (https://feasts.tidyverts.org) to detrend.

## Single-fly sleep and circadian rhythm recording

96-well white Microfluor 2 plates (Fishier) with 400 μl of food (5% sucrose and 1% agar) were loaded with adult male flies (aged 3–5 days). Flies were entrained to the 12Lh:12 h LD cycles for four days at 25 °C to record sleep behavior, then changed to constant darkness for 5-6 days to record circadian rhythms in the absence of light inputs. The fly movement was monitored using a camera at 10s intervals, and the data were then used by the sleep and circadian analysis program SCAMP to analyze sleep and circadian rhythm^86–88^. It calculates activity by shifting the position of *Drosophila* every 10 seconds and calculates sleep using the standard definition (*Drosophila* is recorded as asleep if it remains motionless for at least 5 minutes). For all sleep experiments in Figures 5, 6, S7, and S9, experimental and control groups were assayed concurrently within the same experimental round to minimize batch effects. Bilateral controls (GAL4 driver alone and UAS effector alone) were included for each experimental genotype to validate specificity. Due to the concurrent nature of these assays, the R54D11-GAL4/+ line was used as a shared control for all experimental groups targeting the GAL4 driver side Sample sizes (at least 6–8 flies per genotype) were determined based on prior studies demonstrating reliable detection of robust phenotypes in bilateral control designs^89,90^.

## Single-nucleus RNA-sequencing analyses

snRNAseq dataset analyzed in this paper is published^91^ and available at the Nextflow pipelines (VSN, https://github.com/vib-singlecell-nf), the availability of raw and processed datasets for users to explore, and the development of a crowd-annotation platform with voting, comments, and references through SCope (https://flycellatlas.org/scope), linked to an online analysis platform in ASAP (https://asap.epfl.ch/fca). Single-cell RNA sequencing (scRNA-seq) data from the *Drosophila melanogaster* were obtained from the Fly Cell Atlas website (https://flycellatlas.org/scope). The Seurat (v4.2.2) package (https://satijalab.org/seurat) was utilized for data analysis. Violin plots were generated using the “Vlnplot” function, the cell types are split by FCA.

## Gene expression analysis

Single-cell RNA sequencing (scRNA-seq) data from the *Drosophila melanogaster* were obtained from the GEO under the accession code GSE157504^92^. The same integration method was applied. Data from six time points and under LD and DD conditions were read and integrated using the integration functions provided by the Seurat 4 (version 4.2.2) package^93^. The UMIs data were retrieved, consisting of a grand total of 4,634 cells. The “NormalizeData” function was utilized for the purpose of automated data normalization. Ultimately, we conducted principal component analysis (PCA) on gene expression vectors that were scaled using z-scores.

Subsequently, we limited the data to include just the top 40 PCA components. We employed the ‘FindNeighbors’ and ‘FindClusters’ functions from the Seurat package to cluster the data that had been decreased in dimensions. We utilized t-distributed stochastic neighbor embedding (t-SNE) to generate a two-dimensional map that displays the clusters. One cluster that highly express the ITP gene were extracted, and then the marker genes were calculated using the Seurat ߢFindMarkers’ function.

## Connectome analysis

Whole brain connectomics data were obtained from FlyWire (https://codex.flywire.ai/)^94–99^. The left ITP-LN_d_ (FlyWire Root ID: 720575940634984800) dataset was used to gather information on the synaptic connections between the presynaptic and the postsynaptic neurons of interest. The connectivity was visualized with Sankey diagram and doughnut diagram by the Plotly R Studio library (https://plotly.com/r/).

## Statistical tests

Statistical analysis of mating duration assay was described previously^9–11^. More than 50 males (naïve, experienced and single) were used for mating duration assay. Our experience suggests that the relative mating duration differences between naïve and experienced condition and singly reared are always consistent; however, both absolute values and the magnitude of the difference in each strain can vary. So, we always include internal controls for each treatment as suggested by previous studies^100^. Therefore, statistical comparisons were made between groups that were naïvely reared, sexually experienced and singly reared within each experiment. As mating duration of males showed normal distribution (Kolmogorov-Smirnov tests, p > 0.05), we used two-sided Student’s t tests. The mean ± standard error (s.e.m) (*\*\*\*\* = p < 0.0001, *** = p < 0.001, ** = p < 0.01, * = p < 0.05*). All analysis was done in GraphPad (Prism). Individual tests and significance are detailed in figure legends. Besides traditional *t*-test for statistical analysis, we added estimation statistics for all MD assays and two group comparing graphs. In short, ‘estimation statistics’ is a simple framework that—while avoiding the pitfalls of significance testing—uses familiar statistical concepts: means, mean differences, and error bars. More importantly, it focuses on the effect size of one’s experiment/intervention, as opposed to significance testing^101^. In comparison to typical NHST plots, estimation graphics have the following five significant advantages such as (1) avoid false dichotomy, (2) display all observed values (3) visualize estimate precision (4) show mean difference distribution. And most importantly (5) by focusing attention on an effect size, the difference diagram encourages quantitative reasoning about the system under study^102^. Thus, we conducted a reanalysis of all our two group data sets using both standard *t* tests and estimate statistics. In 2019, the Society for Neuroscience journal eNeuro instituted a policy recommending the use of estimation graphics as the preferred method for data presentation^103^. For sleep experiments, all statistical analyses were performed using IBM SPSS and Prism software. Data were first tested for normal distribution with the Wilks-Shapiro test. One-way analysis of variance (ANOVA) followed by Tukey-Kramer HSD post hoc tests were applied for multiple group comparisons. The number of animals (n) for each experiment is provided in the figures. Data are presented as mean behavioral responses with error bars indicating the standard error of the mean (SEM), and group differences were considered statistically significant at p < 0.05.

---

**Figure S1.**
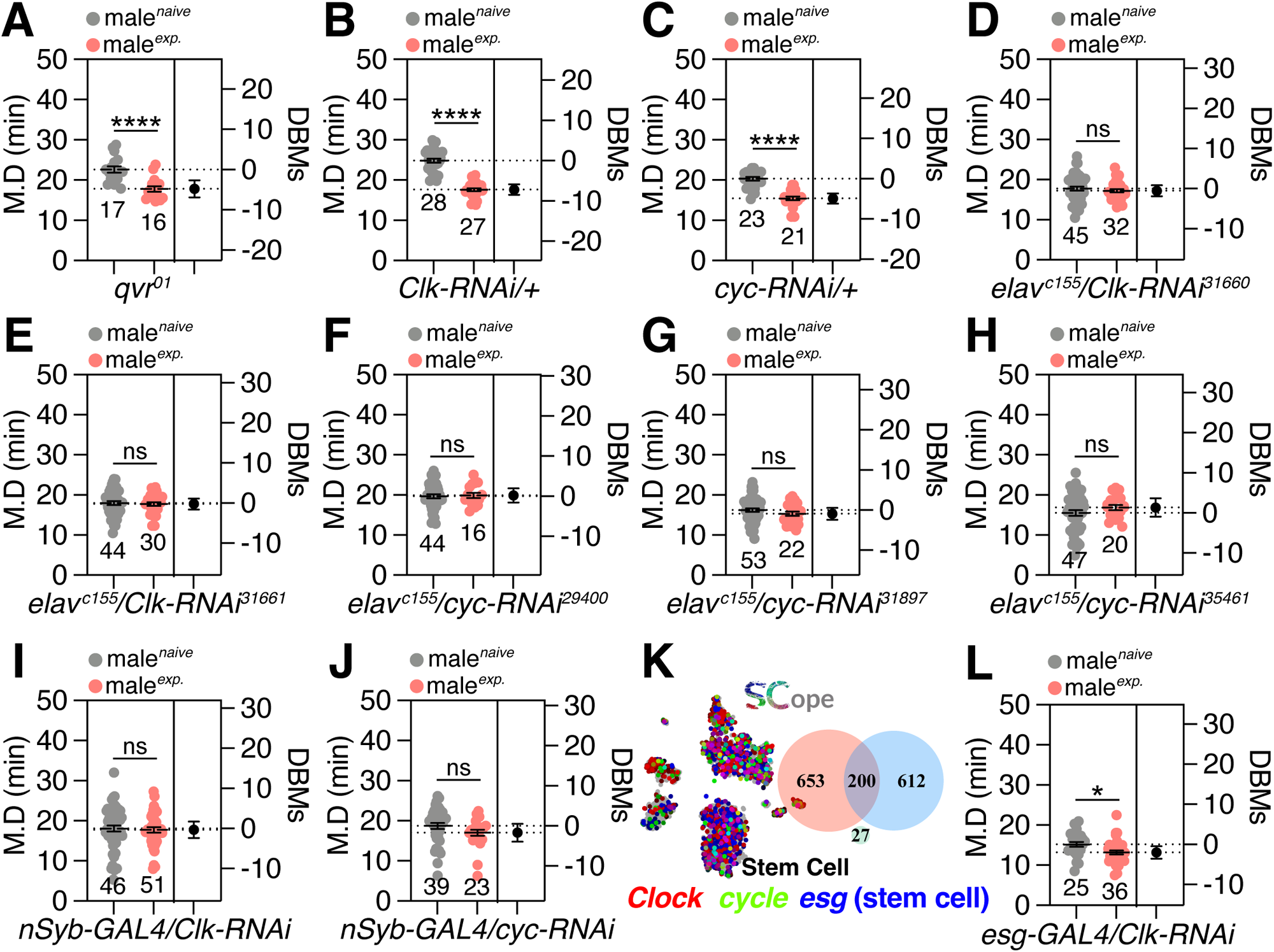
Screening of *Clk-RNAi* lines and *cyc-RNAi* lines. (A) SMD assay of sleepless mutant of *quiver* (*qvr^01^*). (B-C) Genetic control of the SMD assay, which only has (B) *Clk-RNAi* or (C) *cyc-RNAi* with *Canton S* without a GAL4 driver. (D-E) SMD assay for screening of *Clk-RNAi* lines. (F-H) SMD assay for screening of *cyc-RNAi* lines. (I-J) SMD assays for *nSyb-GAL4*-mediated neuronal knockdown of *Clk/cyc* via (I) *Clk-RNAi* and (J) *cyc-RNAi*. (K-L) SCope data of *Clk/cyc* expression in the ‘Stem cell’ population and SMD assays for stem cell-specific knockdown of *Clk* via *Clk-RNAi* using the *esg-GAL4*.

**Figure S2.**
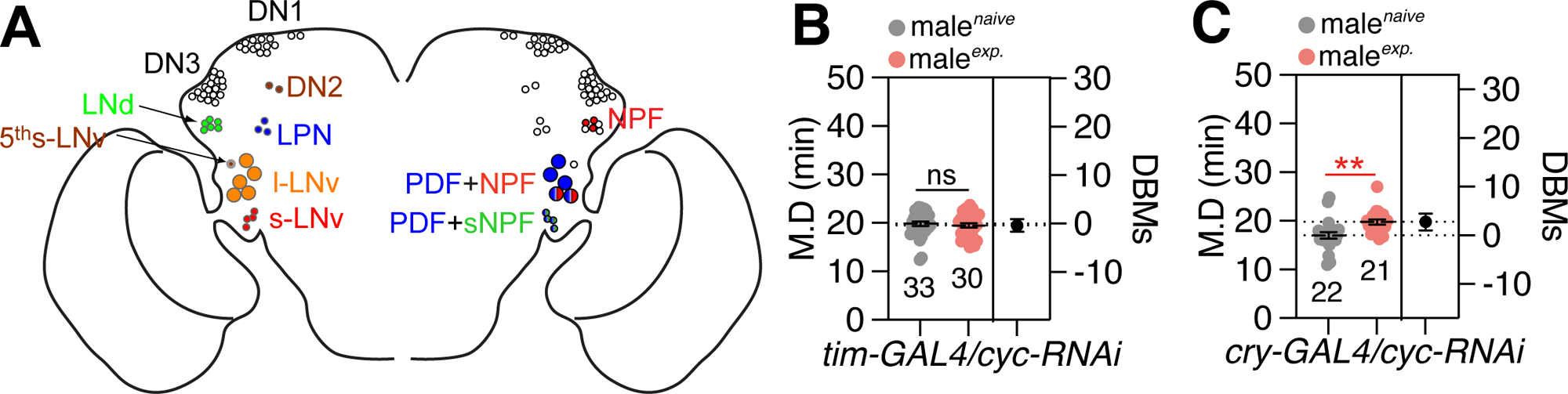
CYC in clock neurons is also critical for interval timing behavior. (A) Clock neurons (left) and their neuropeptide expression (right). (B) SMD assay for *tim-GAL4* driver mediated knockdown of *cyc* in clock neurons via *cyc-RNAi*. (C) SMD assay for *cry-GAL4* driver mediated knockdown of *cyc* in cry-positive neurons via *cyc-RNAi*.

**Figure S3.**
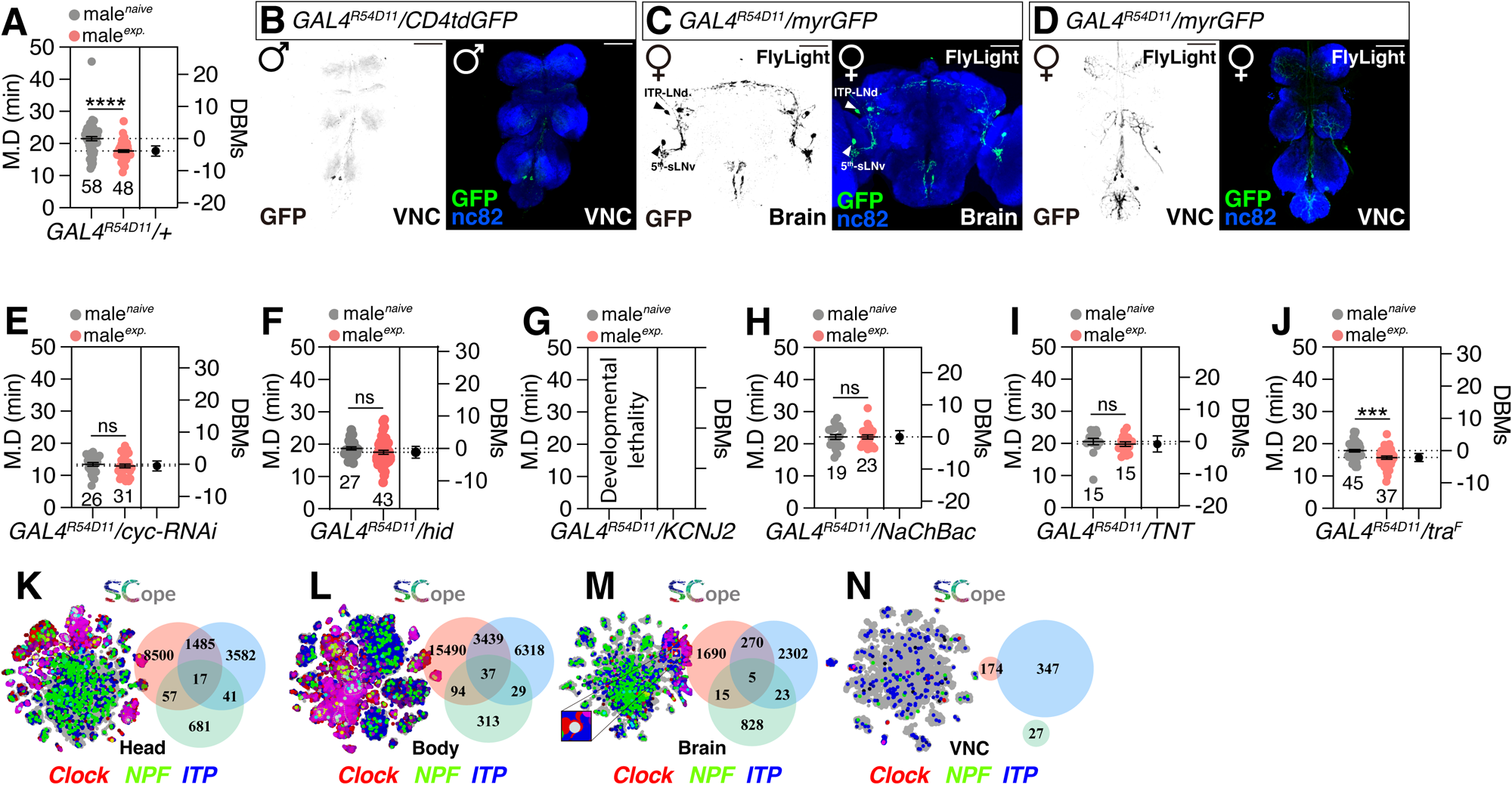
Regular physiological activity of GAL4^R54D11^ neurons is necessary for the formation of interval timing behavior. (A) Genetic control of the SMD assay, which only has *GAL4^R54D11^* with *w^1118^*. (B) VNC of male flies expressing *GAL4^R54D11^* together with *UAS-CD4tdGFP* was immunostained with anti-GFP (green), and nc82 (blue) antibodies. Scale bars represent 50 μm. (C-D) Brain and VNC of female flies expressing *GAL4^R54D11^* together with *UAS-myrGFP* in Flylight. Scale bars represent 100 μm. (E) SMD assay for *GAL4^R54D11^* driver mediated knockdown of *cyc* via *cyc-RNAi*. (F) SMD assay for *GAL4^R54D11^* driver mediated apoptosis in GAL4^R54D11^ neurons via *UAS-hid*. (G) SMD assay for *GAL4^R54D11^* driver mediated electrical silencing in GAL4^R54D11^ neurons via *UAS-KCNJ2.* Flies died during the pupal stage. (H) SMD assay for *GAL4^R54D11^* driver mediated electrical activation in GAL4^R54D11^ neurons via *UAS-NaChBac.* (I) SMD assay for *GAL4^R54D11^*driver mediated inactivation of synaptic transmission in GAL4^R54D11^ neurons via *UAS-TNT.* (J) SMD assay for *GAL4^R54D11^* driver mediated feminization in GAL4^R54D11^ neurons via *UAS-tra^F^.* (K-N) Fly SCope single-cell RNA sequencing data of cells co-expressed Clk together with NPF and ITP in (K) ‘Head’ population, (L) ‘Body’ population, (M) ‘Brain’ population, and (N) ‘VNC’ population.

**Figure S4.**
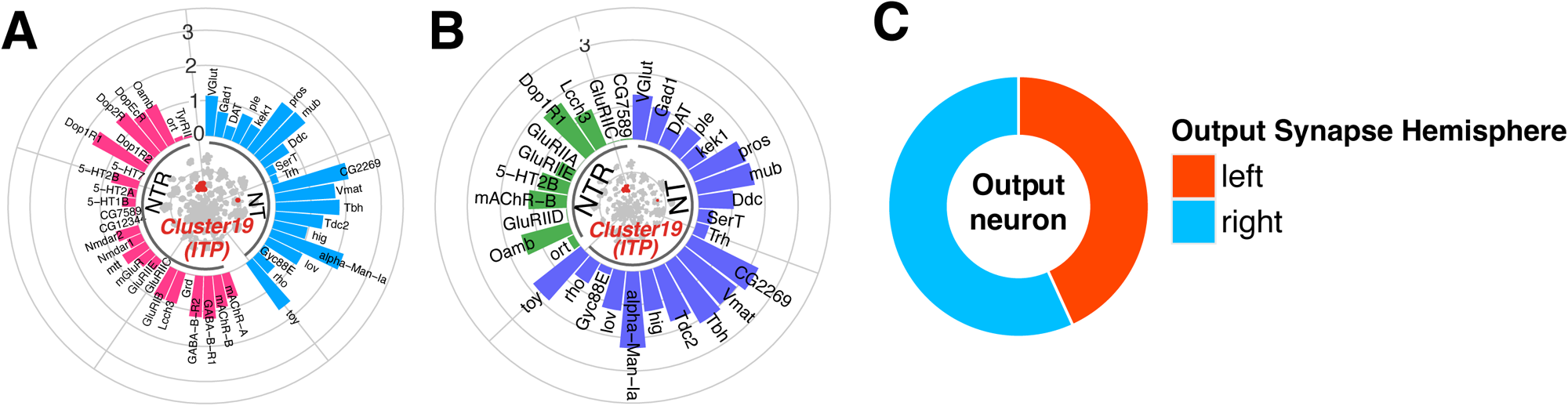
Neurotransmitter inputs and outputs of ITP-LN_d_. (A-B) Expression of neurotransmitter (NT) and neurotransmitter receptor (NTR) in LN_ITP cells. The average UMI count was calculated for each gene in this population, followed by log10 transformation for data normalization. All NT and NTR (A) or genes exhibiting increased expression of NT and NTR (B) were included. (C) Distribution of ITP-LN_d_ output neurons in the left and right brain.

**Figure S5.**
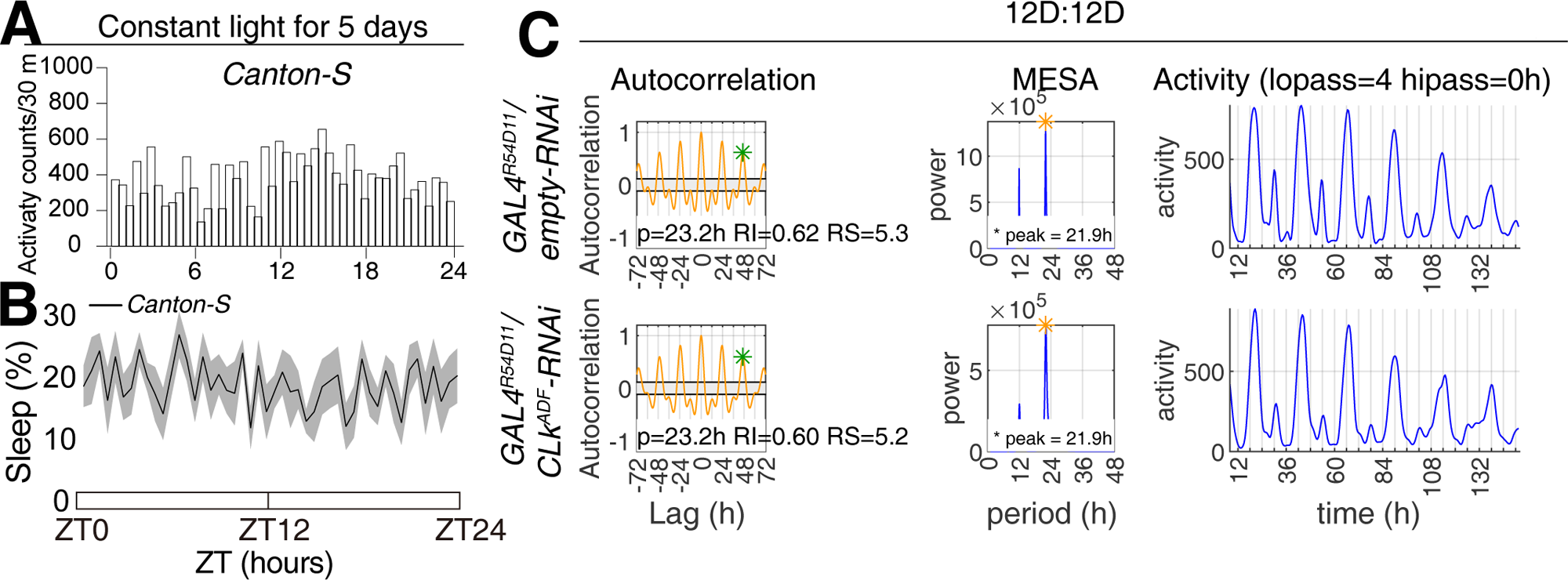
Circadian rhythm of different CLK protein shear forms. (A-B) Activity and sleep profiles of *Canton-S* males under 5 days of constant light. (C) Supplementary data of circadian rhythm for *GAL4^R54D11^* driver mediated knockdown of *Clk-RA*, *RD*, and *RF* via *Clk^ADF^-RNAi*.

**Figure S6.**
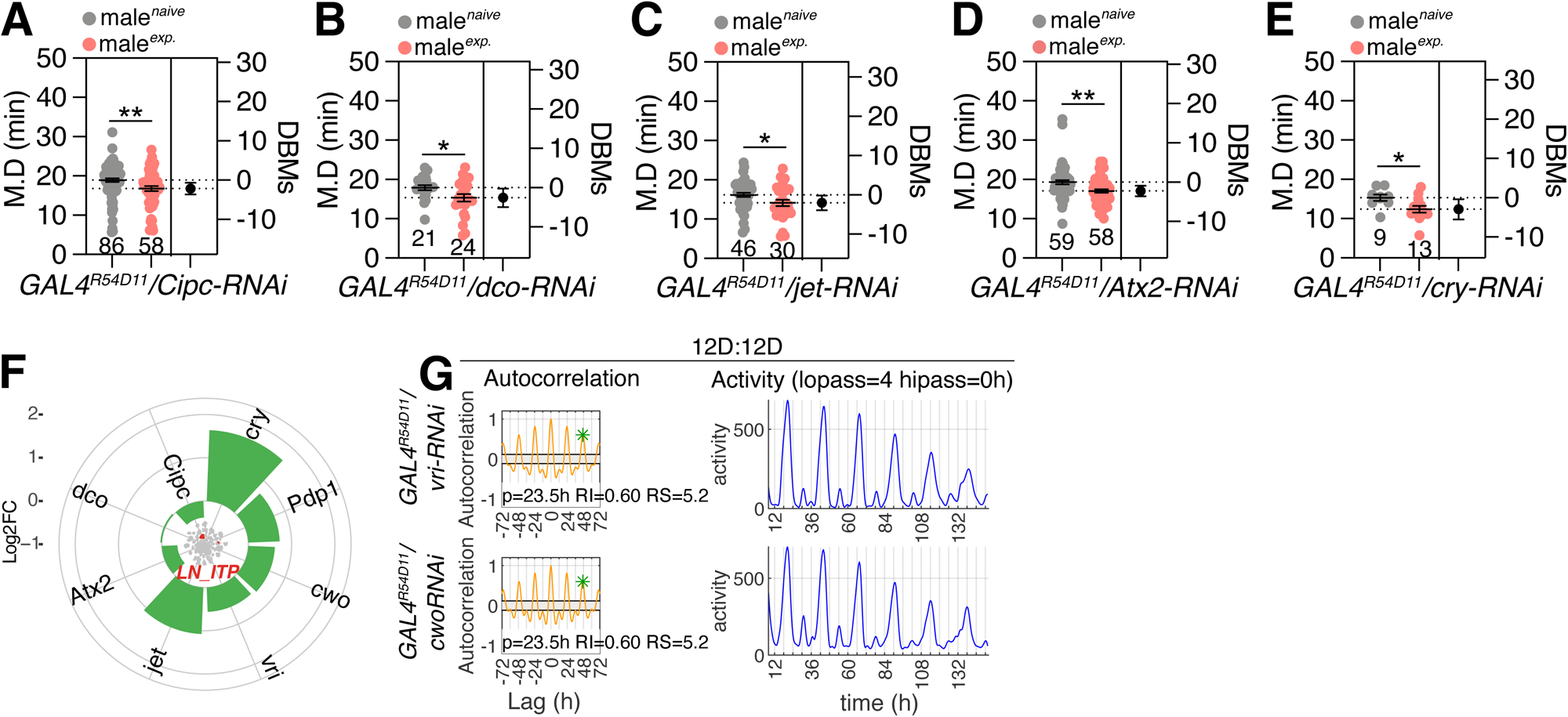
PER/TIM-related genes and some CLK/CYC-associated genes are not involved in regulating interval timing behavior. (A) SMD assay for *GAL4^R54D11^*driver mediated knockdown of CLK/CYC-associated genes via *Cipc-RNAi.* (B-E) SMD assay for *GAL4^R54D11^* driver mediated knockdown of PER/TIM-associated genes via (B) *dco-RNAi*, (C) *jet-RNAi*, (D) *Atx2-RNAi*, and (E) *cry-RNAi*. (F) Log2 Fold Change (Log2FC) of *cry*, *Pdp1*, *cwo*, *vri*, *jet*, *Atx2*, *dco*, and *Cipc* expression in LN_ITP cells versus other brain cells. The Log2FC was calculated using the ‘FindMarkers’ function of the Seurat (v4.2.2) package (https://doi.org/10.1016/j.cell.2021.04.048). (G) Supplementary data of circadian rhythm for *GAL4^R54D11^* driver mediated knockdown of *vri* or *cwo* via *cwo-RNAi* (top) or *vri-RNAi* (bottom).

**Figure S7.**
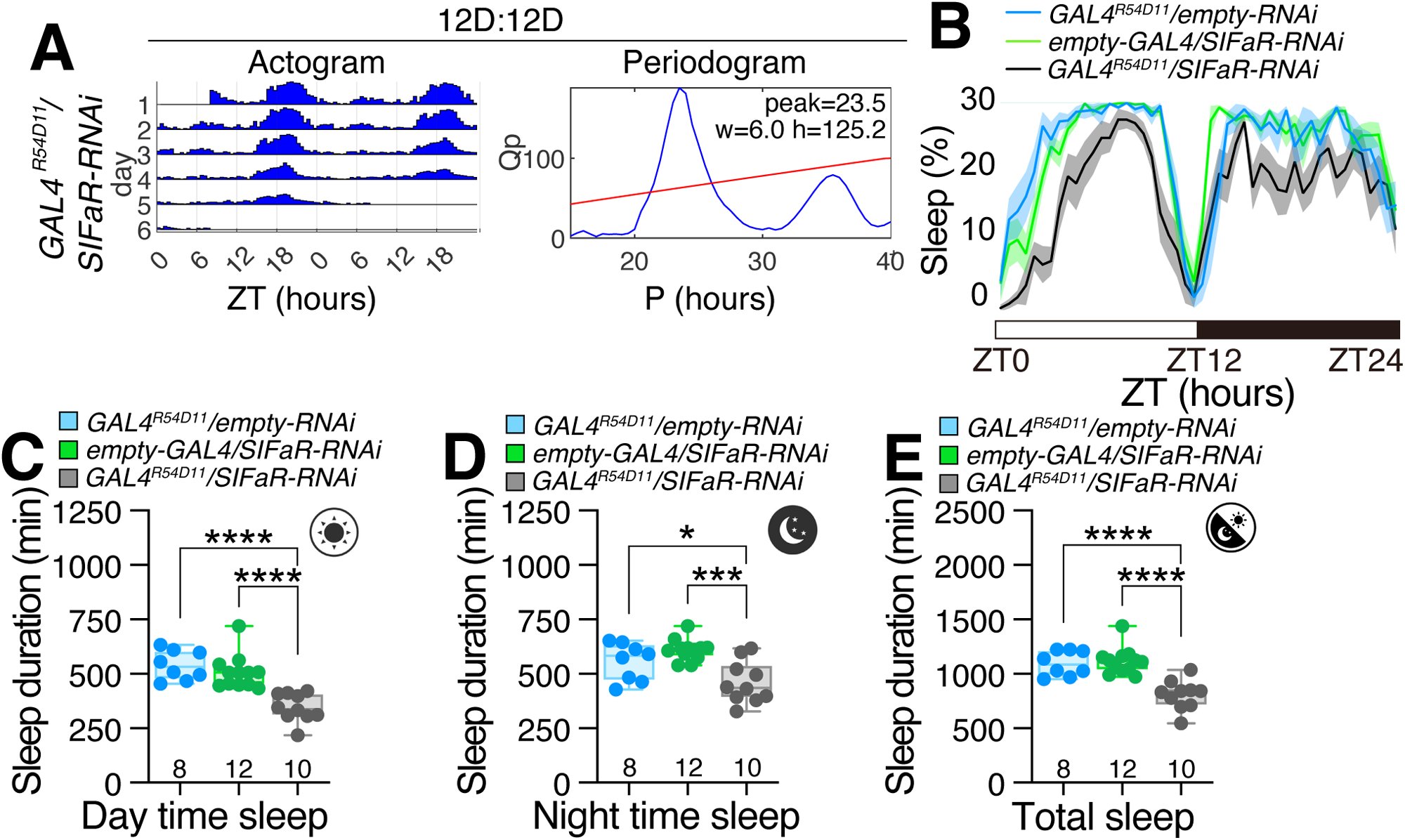
CLK/CYC-associated genes, not SIFaR, are regulating sleep behavior. (A) Actogram and Periodogram of single male flies for *GAL4^R54D11^* driver mediated knockdown of *SIFaR* via *SIFaR-RNAi*. (B-E) Sleep profiles and quantification of flies for *GAL4^R54D11^* mediated knockdown of *SIFaR* via *SIFaR-RNAi*.

**Figure S8.**
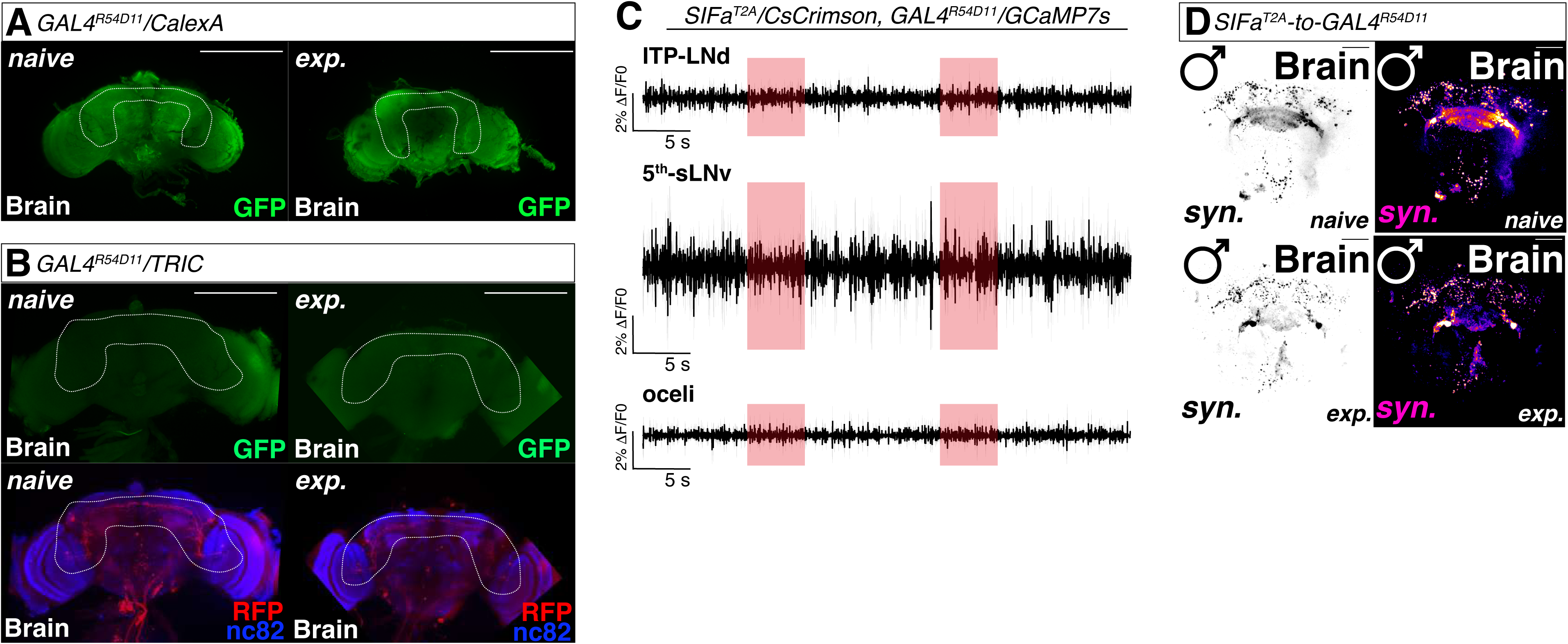
Sexual experiences minimize the SIFa peptidergic signals’ inputs to ITP-LN_d_ neurons without significantly altering the calcium flux. (A) CalexA assay for *GAL4^R54D11^*together with *lexAop-mCD8GFP;UAS-CaLexA, lexAop-CD2-GFP* of naïve (left) and experienced (right) male flies. The dashed line indicates ITP-LN_d_ projection region. Scale bars represent 50 μm. (B) TRIC assay for *GAL4^R54D11^* together with *UAS-IVS-mCD8RFP, LexAop2-mCD8GFP; nSyb-MKII::nlsLexADBDo; UAS-p65AD::CaM* of naïve (left) and experienced (right) male flies. The dashed line indicates ITP-LN_d_ projection region. Scale bars represent 50 μm. (C) GCaMP assay for *SIFa^2A-lexA^* and *GAL4^R54D11^* together with *lexAop-CsChrimson, UAS-GCaMP7s* (ΔF/F0L±LSEM). The red square indicates the time when the red light is turned on. See the “Methods” for a detailed description of the GCaMP experiment in this study. (D) GRASP assay for *SIFa^2A-lexA^* and *GAL4^R54D11^*together with *lexAop-nsyb-spGFP^1-10^, UAS-CD4-spGFP^11^* in SLP and SMP regions of naïve (top) and experienced (bottom) male flies. Scale bars represent 10 μm. Brains of male fly were immunostained with anti-GFP (green) and anti-nc82 (blue) antibodies. The left and bottom panels are presented as a gray scale to clearly show the synapses connection between *SIFa^2A-lexA^* and *GAL4^R54D11^*. GFP is pseudo-colored as “red hot”.

**Figure S9.**
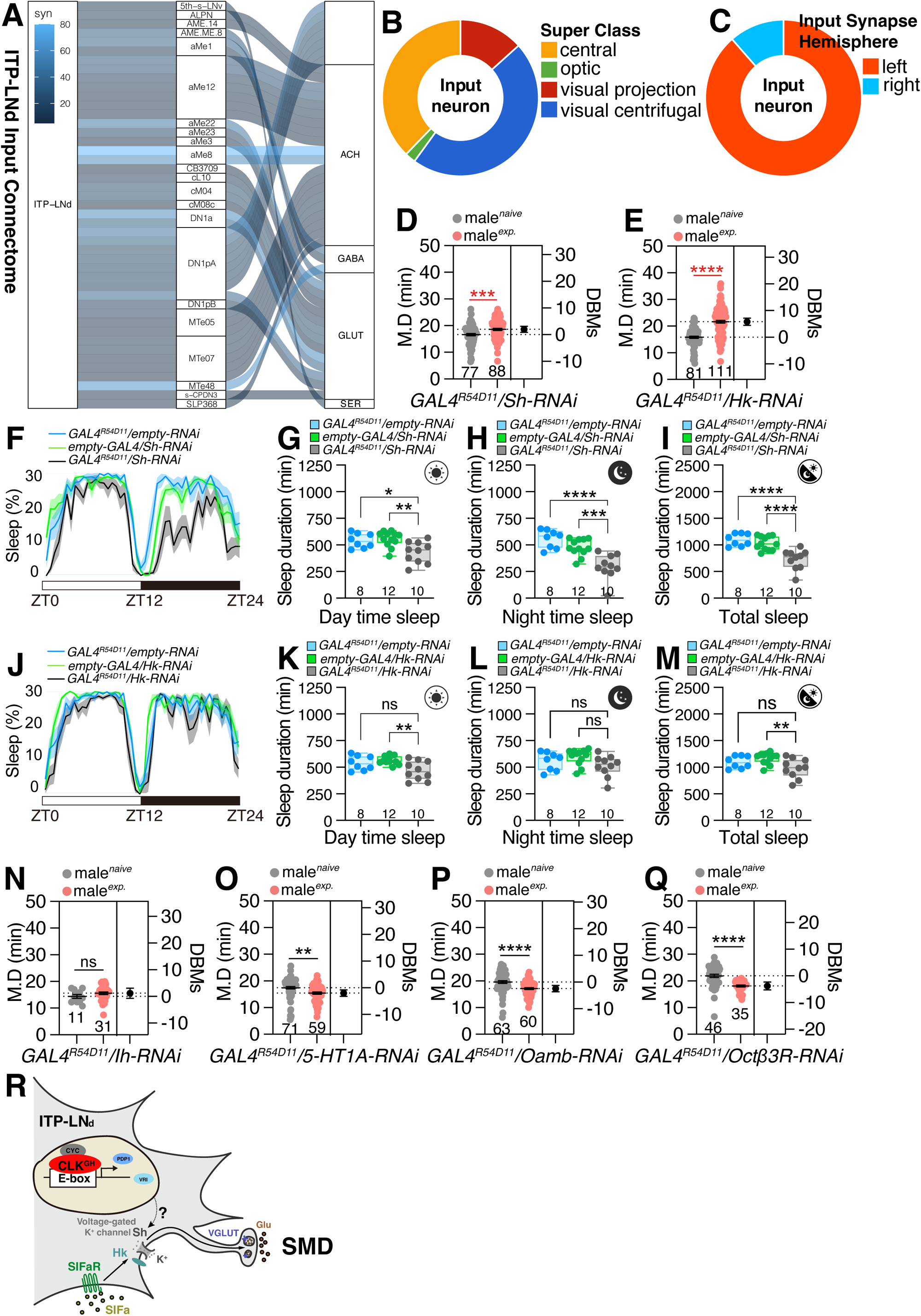
Input of chemical signals and output of electrical signaling in ITP-LN_d_. (A-C) Connectome information of ITP-LN_d_ input neurons. (A) displays a Sankey histogram depicting the synaptic connections between ITP-LN_d_ and input neurons, the neurotransmitters that the input neurons use are displayed on the right. The color indicates the number of synapses connected. (B) displays a donut diagram demonstrating the type of input neurons. (C) displays a donut diagram demonstrating the distribution of ITP-LN_d_ input neurons in the left and right brain. The data is from https://codex.flywire.ai/. (D-E) SMD assay for *GAL4^R54D11^* driver mediated knockdown of ion channels via (D) *Sh-RNAi* and (E) *Hk-RNAi.* (F-M) Sleep profiles and quantification of flies for *GAL4^R54D11^* mediated knockdown of ion channels via (F-I) *Sh-RNAi* and (J-M) *Hk-RNAi.* (N) SMD assay for *GAL4^R54D11^* driver mediated knockdown of ion channel via *Ih-RNAi.* (O-Q) SMD assay for *GAL4^R54D11^* driver mediated knockdown of neurotransmitter receptors via (O) *5-HT1A-RNAi*, (P) *Oamb-RNAi*, and (Q) *Oct*β*3R-RNAi.* (R) Model of how CLK/CYC proteins regulate SMD behavior in ITP-LN_d_. Upon receiving the neuropeptide SIFa from upstream SIFa-positive neurons, ITP-LN_d_ releases Glu to downstream neurons to initiate SMD behavior. CLK^GH^/CYC controls the transmission from neuropeptide signals to neurotransmitter signals by regulating the expression of voltage-gated potassium ion channels Sh/Hk.

