## Supplementary material for "CLOCK-dependent pathway in a single pair of LN_d_ neurons instruct circadian-independent interval timing behavior": Table 1. Screen of 150 clock neurons.

Screen of 150 clock neurons

| | Clusters | GAL4s | SMD | Number (naïve/exp.) |
|---|---|---|---|---|
| **DNs** | $DN1_a$ | *R16C05* | **** | 28/28 |
| | $DN1_p$ | *VT027231*<br>*CNMa* | ****<br>**** | 44/45<br>41/39 |
| | DN2 | $Clk^{int1-3}$ | ** | 66/73 |
| | DN3 | *VT027231* | **** | 44/45 |
| $\mathbf{LN_d}$ | $sNPF^+$ $2LN_d$ | *R16C05* | **** | 28/28 |
| | $ITP^+$ $1LN_d$ | *R54D11*<br>*Mai179*<br>*R78G02*<br>*ITP-RC* | n.s.<br>n.s.<br>n.s.<br>n.s. | 43/39<br>35/14<br>33/29<br>58/47 |
| | $NPF^+$ $cry^-$ $2LN_d$ | *npf + cry-GAL80* | ** | 9/18 |
| | Unknown $1LN_d$ | *N/A* | N/A | N/A |
| $\mathbf{LN_v}$ | $l\text{-}LN_v$ | *pdf* | **** | 61/60 |
| | $s\text{-}LN_v$ | *pdf*<br>$Clk^{int1-3}$ | ****<br>** | 61/60<br>66/73 |
| | $5^{th}$ $s\text{-}LN_v$ | *N/A* | N/A | N/A |
| | LPN | *R65D05* | **** | 61/57 |
