## Supplementary figures and images for "CLOCK-dependent pathway in a single pair of LN_d_ neurons instruct circadian-independent interval timing behavior"

### Graphical Abstract

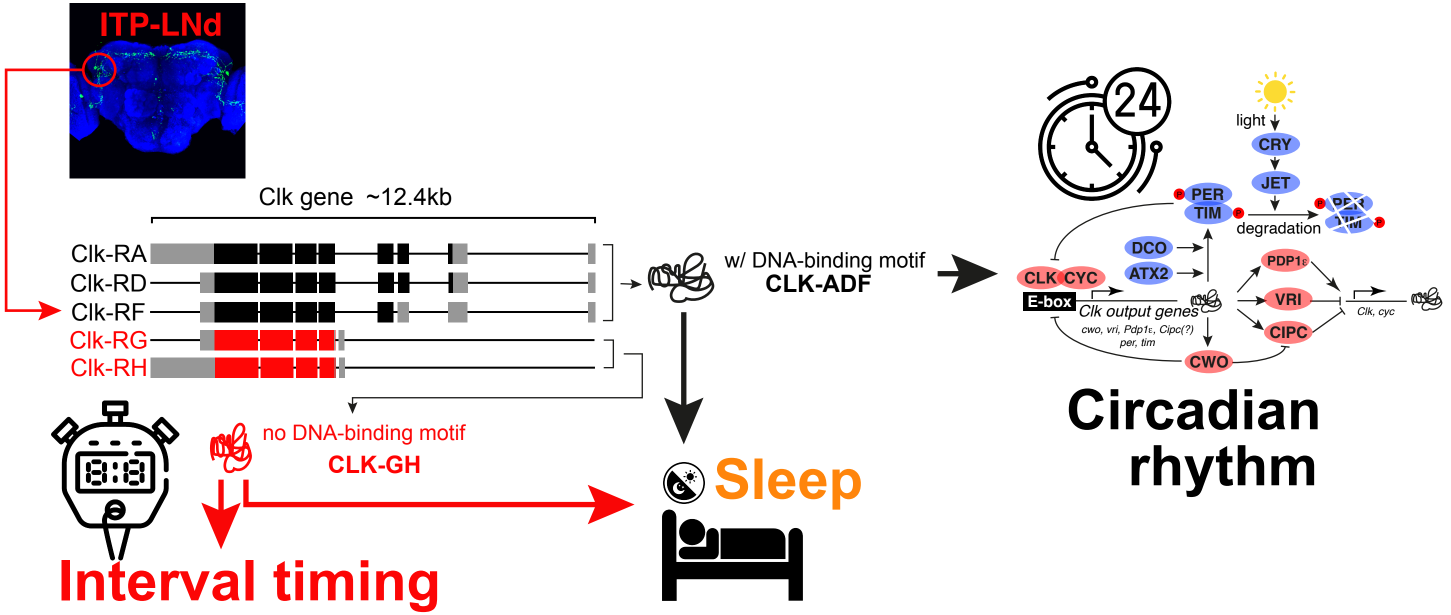


***Graphic Abstract***
